# An Acyclic nucleoside phosphonate effectively blocks the egress of the malaria parasite by inhibiting the synthesis of cyclic GMP

**DOI:** 10.1101/2025.04.01.645218

**Authors:** Marie Ali, Rea Dura, Marc-Antoine Guery, Emma Colard-Itté, Thomas Cheviet, Léa Robresco, Laurence Berry, Corinne Lionne, Catherine Lavazec, Antoine Claessens, Suzanne Peyrottes, Kai Wengelnik, Sharon Wein, Rachel Cerdan

## Abstract

The urgent need for new antimalarial therapies arises from the alarming spread of malaria parasite resistance to existing drugs. A promising candidate, UA2239, an acyclic nucleoside phosphonate with a guanine as nucleobase, demonstrates rapid and irreversible cytotoxic effects on *Plasmodium* parasites, both *in vitro* and in an animal model. It blocks the active exit process, named egress, of merozoites and gametes from infected erythrocytes. UA2239 disrupts the essential cGMP-dependent egress pathway by decreasing cGMP levels in the parasite, making guanylate cyclase (*Pf*GCα) the most likely target. We also uncovered the remarkable molecular mechanism of resistance developed by parasites after prolonged exposure to the drug, which involves mutating not the target itself, but a downstream effector. The unique mechanism action of UA2239 makes it a valuable first-in-class candidate for further development and its ability to inhibit both parasite growth and transmission highlights its therapeutic potential as a dual-stage antimalarial agent.

## Introduction

Malaria is caused by parasites of the genus *Plasmodium* with most fatal cases occurring after *P. falciparum* infection. Despite the effort and progress made in recent decades to control the disease (*1*), the number of deaths due to malaria is still estimated at 597,000 in 2023. After a significant increase at the beginning of the COVID-19 pandemic, malaria cases and deaths have been falling slowly for the last three years, but remain higher than before the pandemic. The very first vaccines are currently administered in a selected number of African countries (*1, 2*) and other vaccine candidates are under development (*3, 4*). Nevertheless, chemotherapy remains indispensable to treat the estimated 263 million cases every year (*1*). Current treatments are based on artemisinin combination therapy (*1*). The emergence of partial resistance to artemisinin in Asia (*5*) and more recently in Africa (*1*), in addition to already established resistance to all commercially available drugs, worsens the situation and emphasizes the urgency of finding alternative options. Effective treatments are therefore necessary to bolster the therapeutic arsenal. The development of new chemical entities with novel mechanisms of action is a priority to face the increasing resistance of parasites to available medicines. Future antimalarial treatments are expected to target at least two stages of the complex life cycle of the parasite, in the human host or during the transmission by mosquitoes (*6*). In humans, *P. falciparum* multiplies initially in liver cells and after release of merozoites into the blood stream, replicates asexually in red blood cells (RBCs) in 48-hour cycles of invasion-growth-multiplication. Within the RBC, the parasite’s asexual cycle progresses through three stages: ring, trophozoite, and schizont. This culminates in the formation of up to 32 daughter cells (merozoites), which actively exit the host cell, a process named egress, to invade new RBCs (*7*). A fraction of parasites differentiates into transmissible sexual forms (male and female gametocytes) by passing through five morphological stages (stage I to V) over 10 to 12 days. After a blood meal, male and female gametocytes ultimately egress and mate in the mosquito gut (*8*). After several weeks of growth and multiplication, parasites settle in the salivary glands ready to be injected in a human host.

Recently, we described a novel series of nucleotide analogs, belonging to the family of acyclic nucleoside phosphonates (ANPs), that show remarkable antimalarial properties (*9*). The best activities were observed with the chemical series of purine analogs (*9*). The lead compound UA2239 (Fig. 1A), a guanosine monophosphate analog has strong activity *in vitro* on *P. falciparum* with a concentration to inhibit 50% of the parasite growth (IC_50_) of 74 nM for the 3D7 strain and of 30 nM and 45 nM for the chloroquine-resistant strains FcM29 and W2, respectively. UA2239 is also active *in vivo* in *P. berghei*-infected mice with a low efficient dose to inhibit 50% of the parasite growth (ED_50_) of 0.5 mg/kg after intraperitoneal administration. UA2239 has no activity on mammalian K562 cells resulting in a very high selectivity index (SI > 10,000) (*9*). Nucleoside- and nucleotide analogs are widely used as therapeutic agents in the clinical treatment of viral infections and of cancer due to their anti-proliferative properties (*10, 11*). Their mechanism of action generally involves their conversion into the corresponding poly-phosphorylated derivatives and their interaction with cellular or viral polymerases (as substrates and/or competitive inhibitors) (*10*). Surprisingly, initial phenotypic characterization of the mode of action of UA2239 on *P. falciparum* revealed no effect on parasite intraerythrocytic development including the formation of merozoites. This contradicts the hypothesis of inhibiting DNA synthesis and suggests a block of merozoite egress (*9*). In *Plasmodium*, both merozoites and gametocytes exit from the RBC in a tightly controlled manner involving several layers of regulation (*12, 13*). Some minutes prior to rupture of the parasitophorous vacuole membrane (PVM), accumulation of the intracellular messenger cyclic GMP (cGMP) leads to the activation of cGMP-dependent protein kinase (PKG) that in turn triggers a calcium signal that is essential for the continuation of the egress process (*14–16*) (Fig. 1B). The cellular level of cGMP is the result of a balanced and tightly regulated activity of its synthesis from GTP by guanylate cyclase (GC) (*17*), and its hydrolysis to GMP by phosphodiesterases (PDEs) (*18*) (Fig. 1B). Specific inhibitors of *Pf*PKG have been developed and lead to impairment of egress with the fully segmented parasites being blocked within the RBC. Compound 2 (C2, also known as ML1, an imidazopyridine derivative) targets the catalytic domain of *Pf*PKG and a medicinal chemistry program led to the development of ML10, a more potent C2 analog (*19–21*). C2 blocks parasite egress reversibly (*14*), and is therefore widely used in *Plasmodium* cell biology. Another compound belonging to the trisubstituted imidazole family also inhibits *Pf*PKG with a potent activity against liver stages, and asexual and sexual blood stages (*22*). In contrast, inhibitors of *Pf*PDEs that increase cellular levels of cGMP induce premature egress of merozoites, impairing the invasion of new RBC (*23–26*). Among this trio of enzymes, PDE, GC and PKG, the only enzyme for which no inhibitor has yet been developed is GC, despite the importance of the cGMP-dependent cellular processes for parasite survival. The *Plasmodium* genome encodes two GC isoforms, the GCα essential at the blood stage and for gametogenesis (*17*) and GCβ required at the mosquito stage for ookinete development (*27*).

**Fig. 1.**
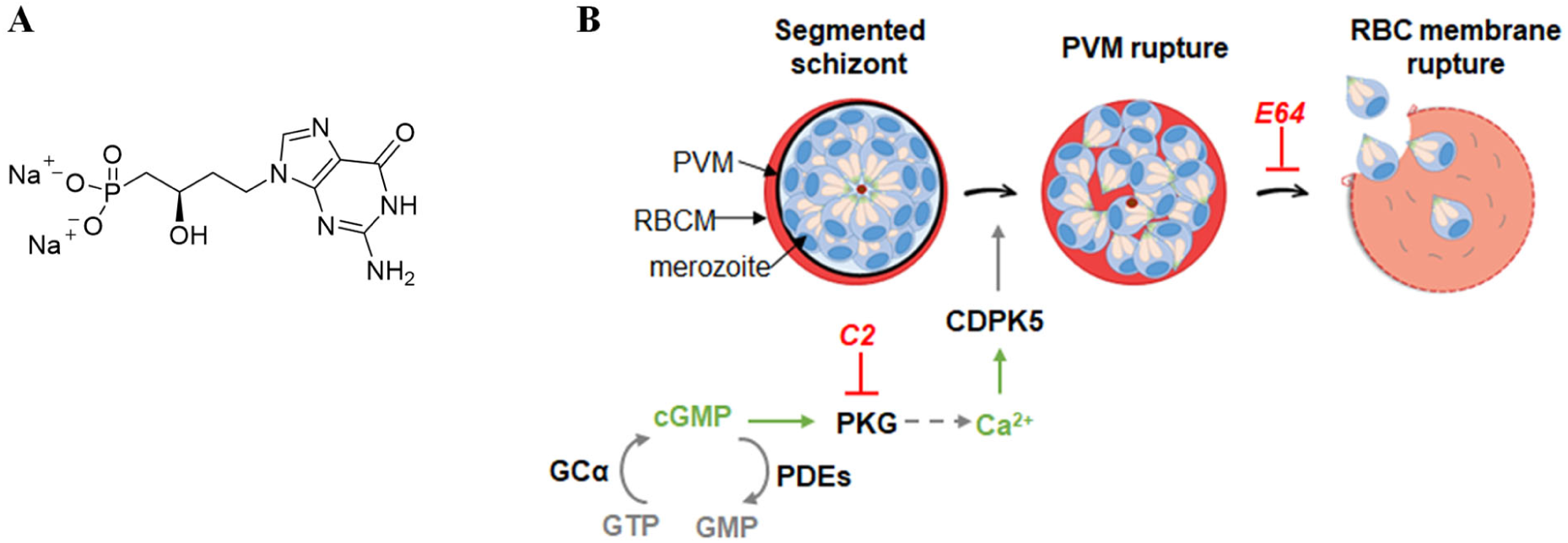
Chemical structure of UA2239 (A) and schematic representation of merozoite egress and the signaling cascades. **(B)**. PVM: parasitophorous vacuole membrane; RBC: red blood cell; RBCM: red blood cell membrane; PKG: cGMP dependent protein kinase; GCα: guanylate cyclase alpha; PDEs: phosphodiesterases; CDPK5: Calcium-Dependent Protein Kinase 5. Inhibitions by C2 and E64 are indicated in red.

In this work, we characterized the pharmacological properties of UA2239 and showed its activity against the symptomatic and transmission stages of the disease. The compound does not inhibit DNA synthesis but irreversibly blocks parasite egress. Its unique mode of action consists of disrupting the crucial cGMP-dependent egress pathway by reducing the level of cGMP in the parasite. We have also discovered the astonishing molecular mechanism of resistance put in place by resistant parasites exposed to prolonged drug treatment.

## Results

### UA2239 is active on trophozoites and schizonts and enters infected red blood cells *via New Permeation Pathways*

We analyzed whether susceptibility to UA2239 differs according to the stage of parasite development. The compound was not active when used over 6 h at the ring stage (Fig. 2A). However, a treatment during trophozoite or schizont development resulted in an IC_50_ of 178 nM and 143 nM, respectively. These values are close to the IC_50_ of 74 nM found when parasites are treated for 48 h. To determine the minimum contact time with the drug required to observe a significant effect, we performed time-course experiments.

**Fig. 2.**
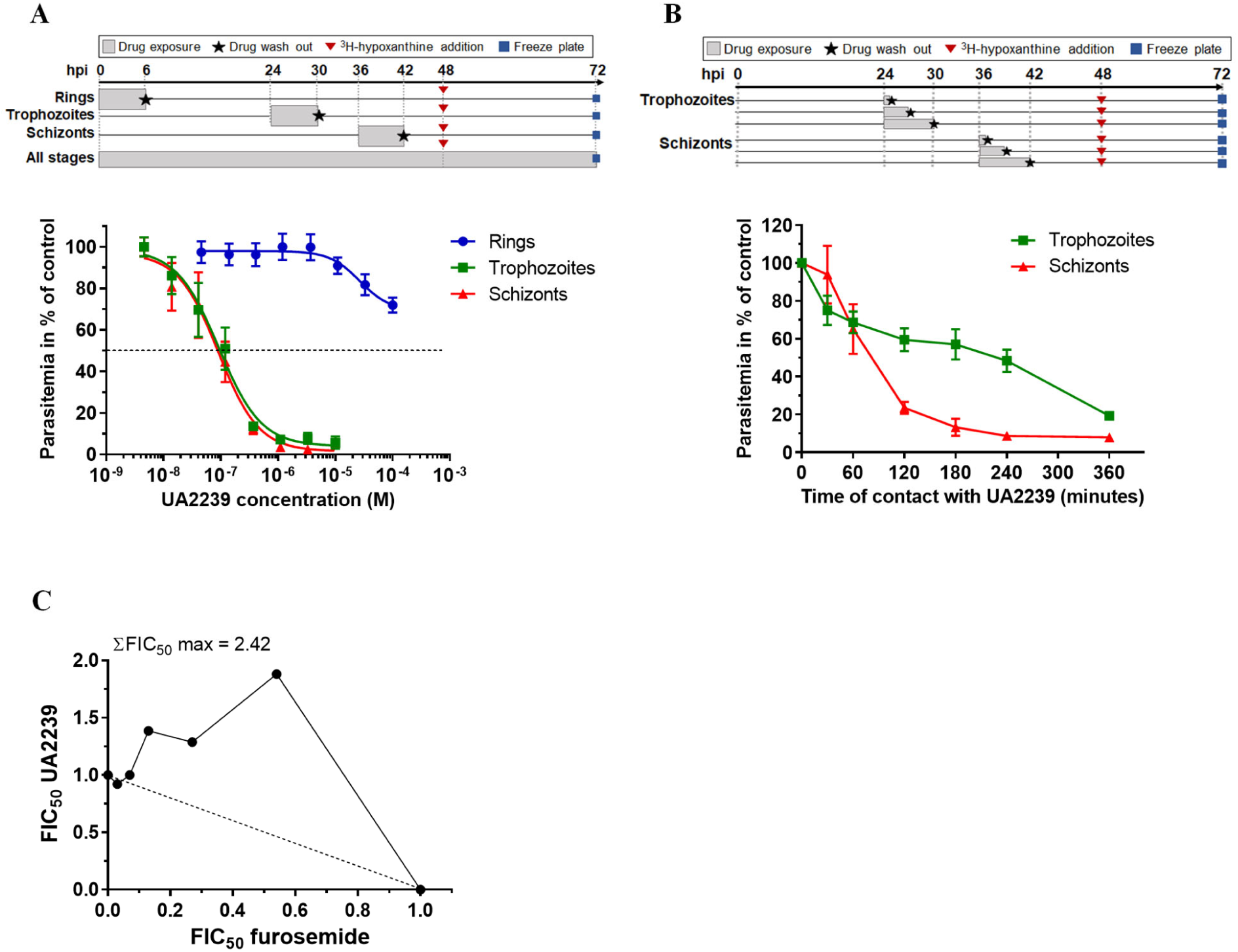
Activity of UA2239 on asexual blood stage parasites. **(A)** Activity of UA2239 on ring, trophozoite and schizont stages after 6 h of contact with various concentrations of UA2239 (n=3, mean ± SEM). The experimental set-up is indicated at the top. **(B)** Time-course of *P. falciparum* growth inhibition by UA2239 dependent on parasite stage. Synchronous cultures at the trophozoite stage, 24 hours post invasion (hpi) or schizont stage, 36 hpi, were treated with 750 nM UA2239 for the indicated time. Cells were then washed and resuspended in fresh media as shown in the scheme at the top (n=3; mean ± SEM). **(C)** Interaction between UA2239 and furosemide represented as an isobologram. Data presented are from one typical experiment (n=3). The maximal sum of FICs of UA2239 and furosemide in combination (ΣFIC50 max) is indicated on the graph.

Synchronized cultures at the trophozoite or schizont stages were pulsed from 0.5 h to 6 h with 750 nM UA2239 (∼10⊆IC_50_) (Fig. 2B). At the schizont stage, a two-hour contact with the compound was sufficient to kill 75% of the parasites and an additional hour of incubation resulted in 90% inhibition of parasite growth. A six-hour contact was required to achieve 75% inhibition growth at the trophozoite stage (Fig. 2B). UA2239 thus exerts a rapid and irreversible cytotoxic effect on both trophozoites and schizonts. This suggests that the compound either acts slowly or remains within the parasite to exert its effect at the end of the cycle. In order to explain the difference observed between early (ring) and late (trophozoite/schizont) stages, we explored the mechanism of entry of UA2239 in the infected RBC (iRBC). One common chemical attribute of all ANPs is their anionic character that limits their passive diffusion across biological membranes. *Plasmodium* relies on specific channels in the iRBC membrane known as new permeation pathways (NPPs) to facilitate the uptake of molecules of low molecular weight such as carbohydrates, nucleosides, nucleobases, ions, etc (*28*). These channels are induced by the parasite and are active in trophozoites and schizonts. To test whether UA2239 enters the iRBC *via* NPPs, we quantified the effect of furosemide, a potent inhibitor of the NPPs (*29*), on UA2239 activity. The isobologram method allows to determine if two compounds in combination exert antagonistic, additive or synergistic effects (*30*). The maximal sum of the fractions of IC_50_ (ΣFIC_50_ max) was found to be 2.25 ± 0.12 (mean ± SEM, n=3) demonstrating that furosemide reduces the antimalarial activity of UA2239 and thus acts as an antagonist (Fig. 2C). This result indicates that UA2239 enters the iRBC through NPPs and is consistent with the lack of UA2239 activity at the ring stage, when NPPs are not yet present.

### UA2239-treated parasites are blocked at the egress step

Since ANPs are described as inhibitors of DNA synthesis (*10, 31–34*), we initially analyzed the effect of UA2239 treatment on parasite DNA content by flow cytometry. For both the control and the treated cultures, the DNA content increased similarly over time from 1 copy (1 N) up to 16 - 32 N showing that DNA synthesis was not affected (Fig. 3A). UA2239-treated parasites developed normally until the late schizont stage and generated merozoites (Fig. 3B) but failed to progress to the next cycle. Merozoite egress is a well characterized process in which the PVM ruptures before lysis of the RBC membrane (Fig. 1B). To identify the specific blockade point in UA2239-treated parasites, we assessed PVM integrity by fluorescence microscopy using a transgenic *P. falciparum* 3D7 strain expressing GFP-tagged parasitophorous vacuolar protein 1 (*Pf*PV1) (*35*) (Fig. 3C). In case of PVM rupture the PV1-GFP fluorescence diffuses into the RBC cytosol. As controls, we used the *Pf*PKG inhibitor C2 and E64 (*36*), a protease inhibitor, which block egress before and after PVM rupture, respectively (Fig. 1B). UA2239 blocked PVM rupture similarly to C2 treatment, thus inhibiting an initial step of merozoite release (Fig. 3C). Finally, we analyzed the ultrastructure of UA2239-treated parasites by electron microscopy. Treatment was started either at 6 or 36 hours post invasion (hpi) and parasites were imaged at 44-46 hpi (Fig. 3D). 89% and 98% of the segmented iRBCs showed an intact PVM after treatment at 6 or 36 hpi, respectively, similar to what we observed for C2 (82%) (table S1). In addition, the distinct contrast between the RBC cytoplasm and the parasitophorous vacuole lumen suggests that the PVM was neither porated nor ruptured in presence of UA2239 (Fig. 3D, images 3 and 4).

**Fig. 3.**
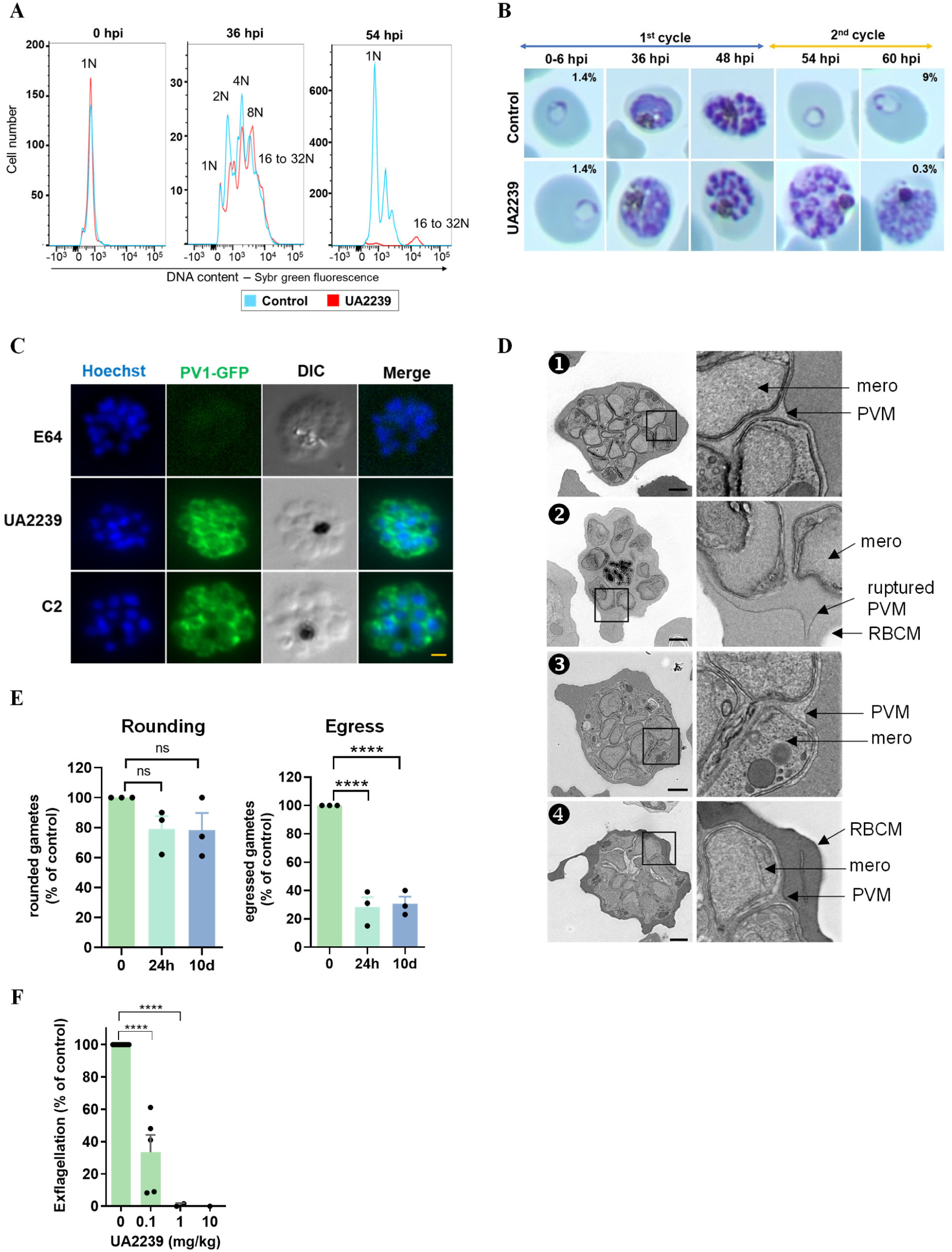
UA2239 inhibits the egress step. **(A)** Effect of UA2239 on DNA content determined by flow cytometry. N represents the genome equivalents. **(B)** Effect of UA2239 on parasite development followed by thin blood smears. Parasitemia is indicated for 6 hpi and 60 hpi. Data are from one of two independent experiments. For (A) and (B), UA2239 concentration was 2 µM. **(C)** IFA on PV1-GFP *P. falciparum* parasites. Late schizont stage parasites were incubated for 4 h with UA2239, C2 or E64. PVM is visualized by the labelling of PV1 protein with GFP. Nuclei were stained with Hoechst (blue). Scale bar 1 µm. (n=3). **(D)** Electron microscopy images of control and UA2239-treated RBCs infected with segmented schizonts. The right panels are close-up views of the framed area on the left panels.1) untreated iRBC with intact PVM, 2) untreated iRBC with a ruptured PVM, 3) iRBCs treated with UA2239 from 6 hpi to 44-46 hpi, 4) iRBCs treated with UA2239 from 36 hpi to 44-46 hpi. mero: merozoite. Scale bar 1 µm. **(E)** Effect of UA2239 (at 10 ⊆ IC_50_) on *P. falciparum* gametogenesis. Gametocytes were treated or not with UA2239 at day 1 for either 24 h or for 10 days. At day 10, percent rounding-up and gamete egress were determined relative to untreated controls (n = 3, mean ± SEM). p values were < 0.0001 using a one-way ANOVA. **(F)** Effect of UA2239 on *P. berghei* gametogenesis after intraperitoneal treatment. No exflagellation center was detected after treatment with 10 mg/kg. (n=2 to 5, mean ± SD). Two-tailed Student’s t-test was used to calculate p value (p<0.0001).

### UA2239 inhibits male gametogenesis of P. falciparum and P. berghei

We investigated whether UA2239 affects the ability of *Plasmodium* parasites to undergo sexual development and gametogenesis. UA2239 was added to synchronous *P. falciparum* gametocyte cultures either daily for 10 days from stage I to stage V, or only during the first 24 h at stage I, because NPPs are mainly active during the first 24 h of gametocyte maturation and then decline in mature gametocyte stages (*37*). In both cases, UA2239 had no effect on gametocytemia, gametocyte morphology and progression of gametocytes from stages I to V (fig. S1). Stage V gametocytes were then subjected to a gametogenesis assay scoring the ability of gametocytes to undergo rounding up and egress from the erythrocyte (*38*). UA2239 had no significant impact on gametocyte rounding up but drastically decreased the proportion of egressed gametes by ∼70% (Fig. 3E). The effect was similar when gametocytes were treated for only 24 h at stage I or for 10 days, consistent with the irreversible activity of UA2239 and the fact that UA2239 enters the infected erythrocyte *via* NPPs.

The effect of UA2239 was also determined *ex vivo* on male gametogenesis of the rodent parasite *P. berghei*. First, exflagellation was quantified in the blood of a *P. berghei* infected mouse and the value was used as positive control. Subsequently, the same mouse was treated with an intraperitoneal dose of UA2239 and exflagellation was again quantified 30 min after treatment. UA2239 reduced the exflagellation of male gametocytes in the mouse model, with an ED_50_ of 0.1 mg/kg (Fig. 3F), a dose of the same order as that observed against the asexual stage (*9*). In summary, our data show that UA2239 inhibits gametogenesis in *P. falciparum* and *P. berghei* at low doses. It is therefore likely that UA2239 blocks sexual reproduction in mosquitoes and hence inhibits parasite transmission.

### UA2239 resistant parasites carry mutations in *Pf*PKG

A common approach to identify the pharmacological target of a compound is to generate resistant parasites and then identify the genetic modifications leading to resistance. We used two different protocols to obtain UA2239-resistant *P. falciparum* blood-stage parasites. Drug pressure was applied intermittently by drug-on/drug-off cycles either using the same concentration (10⊆IC_50_) or gradually increasing the concentration of UA2239 (from 3⊆IC_50_ to 10⊆IC_50_). Parasites were cultured independently and in triplicates for 7 months. From all six independent and resistant populations, several clones were isolated. Two clones per population were selected for whole genome sequencing. Their IC_50_ values for UA2239 had increased at least 30-fold when compared to the IC_50_ of the parental 3D7 strain (74 nM). Focusing on protein-coding sequences, we found non-synonymous mutations in 10 genes (fig. S2A). All clones except one had a single nucleotide polymorphism (SNP) in the *Pfpkg* gene (PF3D7_1436600) resulting in four different and independent substitutions in the *Pf*PKG enzyme (R420I, H524Y, H524N and D597Y) (fig. S2A). *Pf*PKG, one of the central regulators of merozoite egress, is a 853-residue protein, and the C-terminal catalytic domain is preceded by four cGMP-binding (CNB) domains, three of which (A, B, and D) are able to bind cGMP (*16, 39, 40*). Three of the four mutations (R420I, H524N, H524Y) are localized in the CNB-D domain; the fourth mutation (D597Y) is in the catalytic domain (Fig. 4A).

**Fig. 4.**
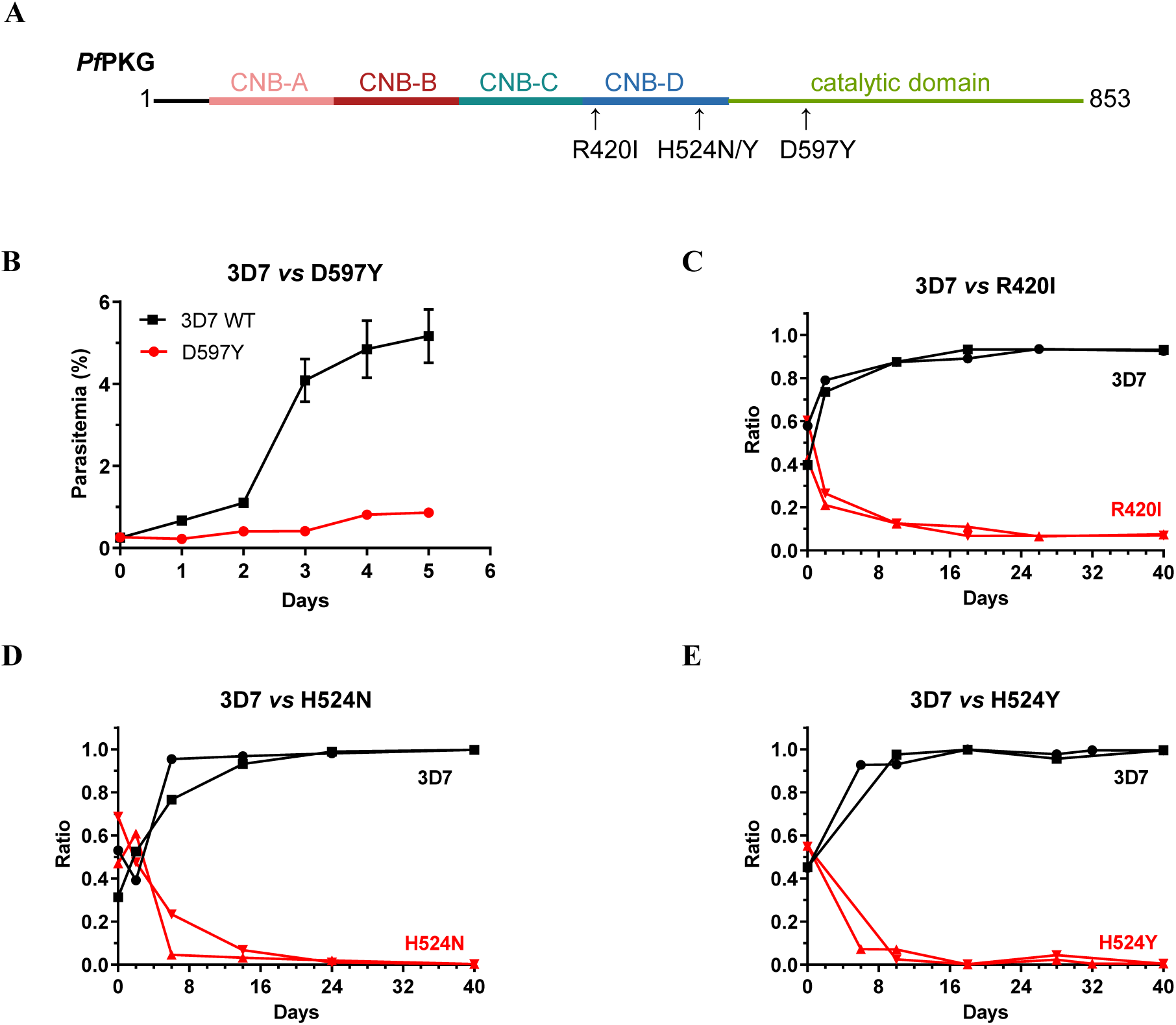
Mutations conferring resistance to UA2239 have a fitness cost for the parasite. **(A)** Schematic representation of the *Pf*PKG domain organization. *Pf*PKG consist of four cyclic nucleotide binding domains (CNB-A, -B, -C and –D) followed by the C-terminal catalytic domain. The positions of the identified mutations are indicated. **(B)** Growth of D597Y GER-parasites was followed by stained thin blood smears over 6 days (n=3). Error bars are within the size of the symbols. **(C-E)** Pairwise competitive growth assays followed by qPCR on genomic DNA for GER-parasites of R420I **(C)**, H524N **(D)** and H524Y **(E)**. Cultures for competition assays were started at approximately equal parasitemia for each strain, equivalent to a ratio of around 0.5 for each strain. This ratio rises to 1 for the fastest-growing strain when it represents 100% of the parasites (n=2).

The second most prevalent mutated gene encodes gametocyte development protein 1 (GDV1, PF3D7_0935400). A stop codon within its reading frame was present in 8 of the 12 clones. GDV1 functions in gametocyte development (*41, 42*) making it unlikely to be directly related to UA2239 resistance. Interestingly, the only clone without a mutation in *Pf*PKG has a non-synonymous mutation in PDEβ (PF3D7_1321500) (fig. S2A). *Pf*PDE acts in the same signaling pathway as *Pf*PKG and negatively regulates the cellular levels of cGMP, the essential activator of *Pf*PKG. Despite the interest of such a mutation for the resistance mechanism, it is highly unlikely that *Pf*PDEβ is the target of UA2239, since the inhibition of *Pf*PDEβ has been described to impair invasion due to premature egress of merozoites (*14, 25, 26*), the opposite of the phenotype observed upon UA2239 treatment. The analysis also revealed 11 micro insertions/deletions (INDELS) in the genomes of the resistant parasites (fig. S2B). None of the corresponding proteins is known to be related to the parasite egress. Copy number variations were not present in the resistant clones.

For further characterization, we selected one clone obtained after drug pressure (termed R-parasites) for each mutation in *Pf*PKG (Table 1 and fig. S2).

**Table 1.** Sensitivity of UA2239-resistant strains (R- and GER-parasites) to UA2239. Data are mean ± SD of 3 independent experiments performed in duplicates except for (**\***) which is the mean of 6 independent experiments performed in duplicates.

| Strain | IC<sub>50</sub> of UA2239 (nM) |  |
| --- | --- | --- |
| WT 3D7 | 74.0 ± 16.2 * |  |
|  | R-parasites | GER-parasites |
| *Pf*PKG - R420I | 2 763 ± 662 | 568 ± 64 |
| *Pf*PKG - H524N | 4 103 ± 1 674 | 1 333 ± 98 |
| *Pf*PKG - H524Y | 2 908 ± 926 | 814 ± 173 |
| *Pf*PKG - D597Y | 3 917 ± 117 | 693 ± 117 |

### Single site mutations of *Pf*PKG confer resistance to UA2239

To confirm the involvement of the mutations in the resistance mechanism, we introduced individually each mutation identified in the 3D7 strain using CRISPR-Cas9 (fig. S3). The four transgenic parasite lines obtained were cloned and genome editing was confirmed by PCR and sequencing (fig. S4). All genome-edited resistant (GER) strains were found to be resistant to UA2239. The IC_50_ values increased 8- to 18-fold with respect to the 3D7 strain (Table 1) demonstrating that each mutation indeed confers resistance to UA2239. It should be noted that the GER-parasites showed lower IC_50_ values than the corresponding R-parasites (Table 1). It is likely that the 7 months *in vitro* culture under drug pressure induced additional adaptations that cannot be detected by genome sequencing.

### Mutations conferring resistance to UA2239 have a fitness cost for the parasite

All GER-parasite clones showed a growth defect. For the PKG-D597Y GER-parasites lower parasitemia was easily observed by counting stained thin blood smears over 6 days. While the multiplication rate of the wild-type strain is about 4 to 5 per cycle, PKG-D597Y showed an average multiplication rate of 1.75 (Fig. 4B). For the other PKG GER-parasites (R420I, H524N or H524Y), the fitness cost was less drastic, allowing to perform competitive growth assays. Equal numbers of synchronous 3D7 and GER-parasites were co-cultured over 20 cycles and their relative abundance was determined by qPCR on genomic DNA. The 3D7 strain outcompeted the genome-edited R420I, H524N or H524Y strains quickly, in less than four cycles (Fig. 4C-E). In conclusion, all mutations conferring resistance to UA2239 induce a heavy fitness cost for the parasite, resulting in a strong growth disadvantage compared to the wild-type strain.

### UA2239-resistant parasites show no cross resistance with standard antimalarial drugs

We determined the sensitivity of GER-parasites to the two standard antimalarial drugs, chloroquine (CQ) and dihydroartemisinin (DHA). None of the *Pf*PKG mutations induced a significant change in the IC_50_ value of CQ compared with the wild-type strain (fig. S5A). For artemisinin, a strain is considered resistant if its survival rate exceeds 1% in a classic ring-stage survival assay (RSA). For the GER-lines, survival rates were <1%, indicating that these mutations do not confer resistance to DHA (Fig. 5A). The same experiments were conducted with the R-parasites and showed similar results (fig. S5A, B).

**Fig. 5.**
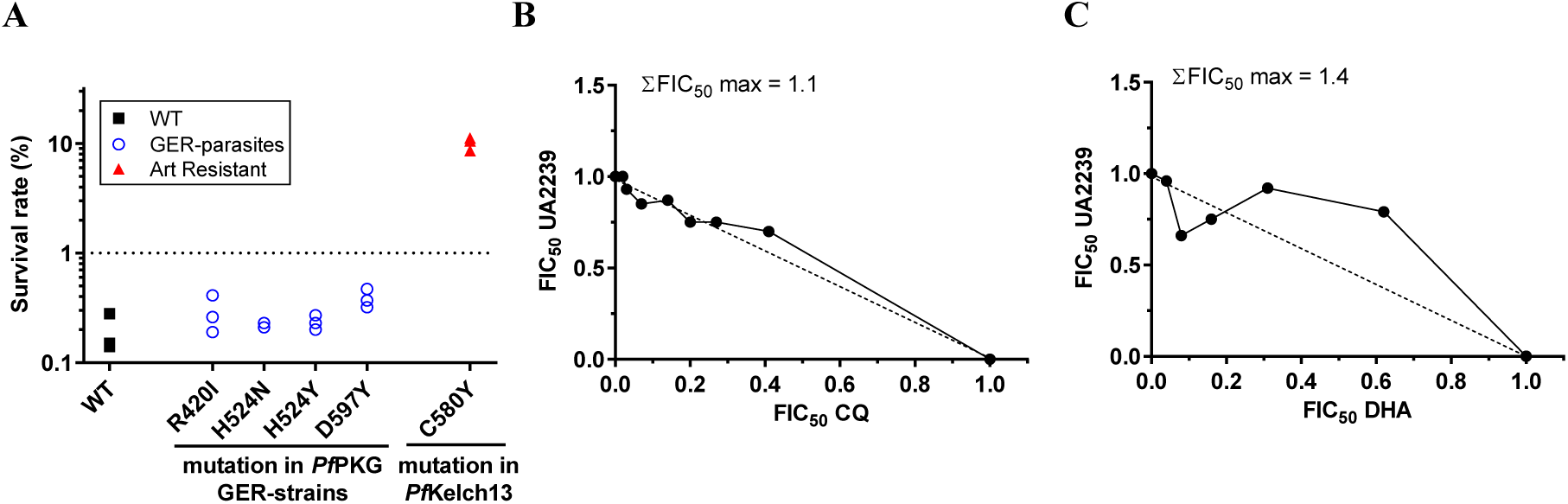
Interaction of UA2239 with other antimalarial drugs. **(A)** Sensitivity of UA2239-resistant strains to dihydroartemisinin (DHA). Sensitivity of GER-parasites to DHA was determined using ring-stage survival assay (RSA). A survival rate <1% indicates sensitive strains (below the dashed line). 3D7 WT strain served as the sensitive control (▪) and the NF54 Kelch13 C580Y line (▴) as resistant control^70^ (n=3 biological replicates). **B-C** *In vitro* interaction between UA2239 and chloroquine (CQ) **(B)** or DHA **(C)** for their antimalarial activities against the 3D7 WT strain. Effect of CQ and DHA on antimalarial activity is represented as an isobologram. Data presented are from one representative experiment (n=3). The maximal ΣFIC_50_ (ΣFIC_50_ max) of the presented experiments are indicated on the graphs.

Intriguingly, only the D597Y R-parasites had a survival rate of around 2% (fig. S5B) while the corresponding D597Y GER-parasites were sensitive to DHA indicating that the D597Y mutation alone cannot be the cause of artemisinin resistance (Fig. 5A).

We also assessed the interactions between UA2239 and CQ or DHA at the pharmacological level using the isobologram method. We observed additive interactions in both combinations, with a mean ΣFIC_50_ max of 1.32±0.15 and 1.34±0.04 (±SEM) for CQ and DHA, respectively (Fig. 5B, C). These results suggest that UA2239 could be used in association with either CQ or DHA.

### UA2239 inhibits the cGMP-PKG egress pathway through a novel mechanism of action

To elucidate the mode of action of UA2239, we first aimed to determine whether PKG is its molecular target. The PKG mutations we identified confer resistance, and UA2239’s effect on parasites resemble those observed with C2. UA2239 could impair *Pf*PKG activity, either by binding to the catalytic site, as C2 does, or by preventing the cGMP-induced activation. To rapidly test the first hypothesis, we took advantage of the availability of both UA2239-resistant and C2-resistant parasites (T618Q) (*43*) that we had previously generated. We assessed the sensitivity of the different R- and GER-parasites to C2 and inversely the sensitivity of a C2-resistant strain to UA2239. Our results revealed the absence of cross-resistance between UA2239 and C2 (Table 2) and an additive interaction between the two compounds (fig. S6). In parallel, we measured the activity of the recombinant full-length *Pf*PKG (provided by the Protein Biochemistry Platform of Geneva University) in the presence of UA2239 or C2 as a control. As expected, *Pf*PKG was activated by cGMP, and this activity was inhibited by C2. Unexpectedly, UA2239 did not impair *Pf*PKG activity, even at a concentration 10 times higher than the one of cGMP (Fig. 6A). Taken together, these results indicate that UA2239 binds neither to the catalytic site nor to the CNB domains of the enzyme, and therefore suggest a different primary target for UA2239 than *Pf*PKG. Yet, single mutations introduced in *Pfpkg* confer resistance to UA2239, suggesting a functional link between UA2239 mechanism of action and the kinase. This enzyme is activated through the cooperative binding of cGMP, whose level is regulated by the balance between its synthesis by GC and its degradation by PDE. Hence, we quantified the intracellular cGMP level and observed a dose-dependent decrease in cGMP content upon UA2239 addition to segmented schizonts (Fig. 6B). When UA2239-treated parasites were exposed to a membrane-permeable cGMP analog (PET-cGMP) able to activate *Pf*PKG (*17, 44*), we observed a significant decrease in the number of segmented parasites and a high increase in the numbers of rings (Fig. 6C) indicating a resumption of parasite growth. The effect of UA2239 can thus be complemented by the addition of the cGMP analog. As expected, PET-cGMP did not complement C2-treated parasites (Fig. 6C).

**Fig. 6.**
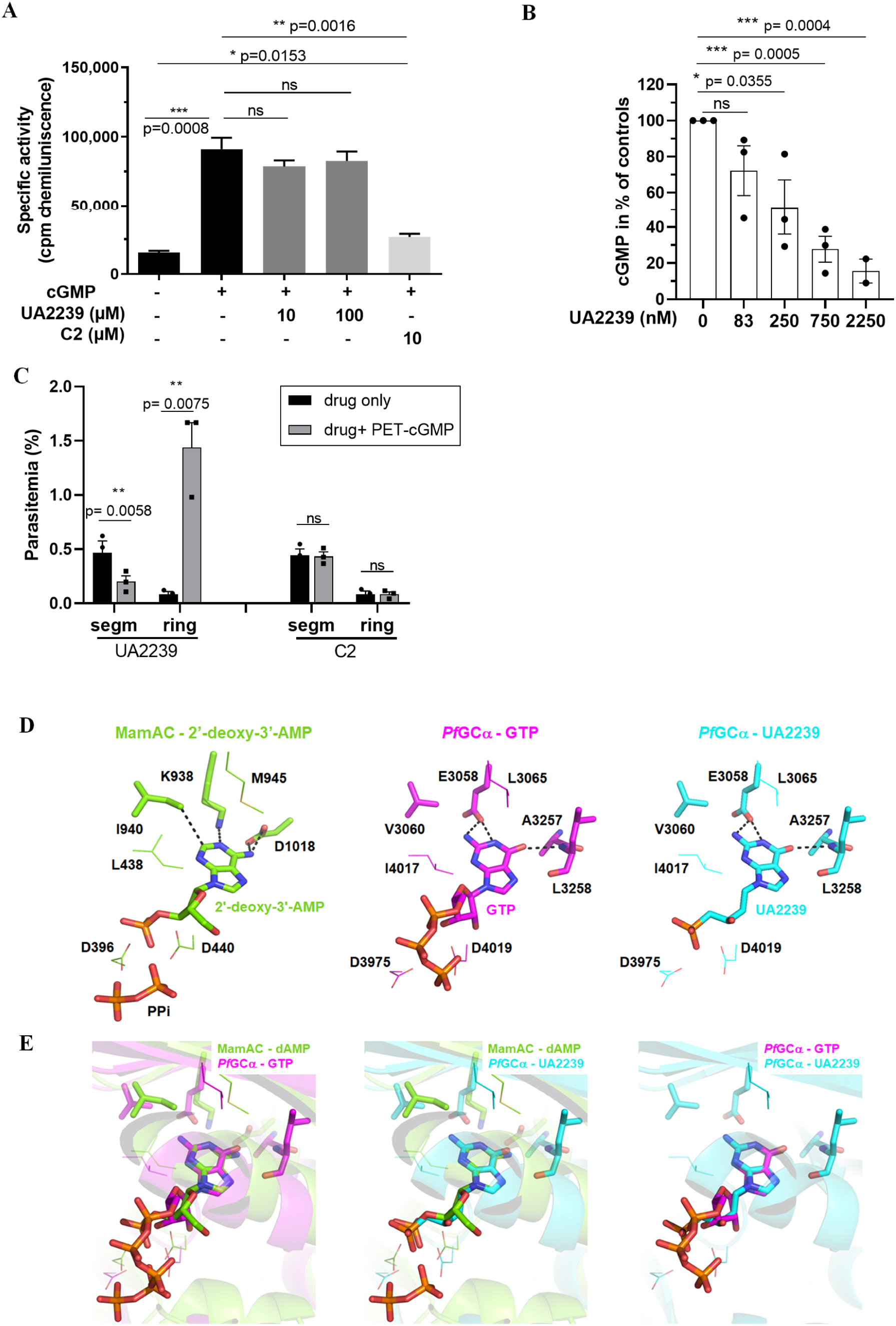
Mechanism of action of UA2239 on cGMP-dependent egress pathway. **(A)** Specific activity of the recombinant full-length *Pf*PKG was determined in presence or absence of 10 µM cGMP using ADP-Glo kinase kit. A final concentration of 100 nM *Pf*PKG was used. UA2239 was added at 10 or 100 µM and C2 at 10 µM. Data presented are from one typical experiment (mean ± SEM). Experiments were done 3 times in triplicate. Two-tailed Student’s t-test was used to calculate p value. **(B)** Effect of UA2239 on intracellular cGMP levels. Purified schizonts (3D7 WT) were allowed to develop in the presence of 1.5 µM C2 (to prevent egress) either with or without various concentrations of UA2239. Segmented schizonts were lysed with 0.1M HCl and cGMP levels were determined. Data are expressed in % of the untreated samples (n=3 independent experiments, mean ± SEM). Two-tailed Student’s t-test was used to calculate p value. **(C)** Effect of PET-cGMP on reversal of parasite egress block and induction of ring-stage development. Ratio paired Students’s t-test was used to calculate p value. **(D)** Binding pockets of MamAC in complex with 2’-deoxy-3’-AMP and PPi (PBD 1CS4) (left panel), of the *Pf*GCα model in complex with the substrate GTP (middle panel) or with UA2239 (right panel). **(E)** Overlays of MamAC (green) and *Pf*GCα (pink GTP ligand or blue UA2239 ligand) (left and middle panels) and overlay of *Pf*GCα with GTP or UA2239 (right panel). The ligands are shown as sticks. The residues sharing conserved interactions with the ligands in the two binding sites are shown as lines. The residues allowing the selective interactions with either an adenine or a guanine base are shown as sticks.

**Table 2.** Sensitivity of UA2239-resistant strains (R- and GER-parasites) to C2 and of C2-resistant strain (T618Q) to UA2239. Data are mean ± SD of 3 independent experiments performed in duplicates except for (**\***) which is the mean of 6 independent experiments performed in duplicates.

| Strain |  | Compound | IC<sub>50</sub> (nM) |  |
| --- | --- | --- | --- | --- |
| WT 3D7 |  | UA2239 | 74.0 ± 16.2* |  |
|  |  | C2 | 284.2 ± 48.2 |  |
|  |  |  | R-parasites | GER-parasites |
| UA2239-resistant | *Pf*PKG -R420I | C2 | 271.4 ± 57.4 | 282.8 ± 60.3 |
|  | *Pf*PKG -H524N | C2 | 128.9 ± 56.8 | 167.4 ± 37.5 |
|  | *Pf*PKG -H524Y | C2 | 116.9 ± 7.1 | 167.4 ± 38.5 |
|  | *Pf*PKG -D597Y | C2 | 128.9 ± 59.9 | 274.2 ± 6.5 |
| C2-resistant | *Pf*PKG -T618Q | UA2239 | - | 145.8 ± 37.4 |
|  | *Pf*PKG -T618Q | C2 | - | 3 511.7 ± 962.0 |

Altogether these results demonstrate that the mechanism of action of UA2239 is due to an insufficient level of cGMP in the parasites, which blocks the signaling cascade at the very beginning (Fig. 1B). Inhibition of cGMP synthesis makes *Pf*GCα the likely primary target.

### UA2239 is docked in the substrate/product binding site of the *Pf*GCα 3D model

In the absence of a known 3D structure, we modelled the guanylate cyclase domain of *Pf*GCα using AlphaFold2 (*45*) to predict whether UA2239 can bind to its active site. *Pf*GCα (PF3D7_1138400) is a bifunctional transmembrane enzyme of 4226 amino acids with an N-terminal P4-ATPase-like domain and a C-terminal GC domain. The GC domain consists of two catalytic sub-domains, C1 and C2, each preceded by six transmembrane helices (*17*). The overall modelled structure is shown in figure S7, where it is compared with the X-ray crystal structure of the cyclase domain of the mammalian adenylate cyclase, MamAC (PDB 1CS4). We chose this protein as a model anchor because, despite its relatively low sequence identity with GC domain of *Pf*GCα (26%), it shares one of the highest identities amongst the protein family and its structure is available at high resolution (<3 Å) and in complex with a substrate analog, the 2’-deoxy-adenosine 3’-monophosphate (2’-deoxy-3’-AMP). The two overall structures align well, with a root-mean-square deviation of 1.86 Å (fig. S7D). GTP, the substrate of *Pf*GCα, and UA2239 were docked into the AlphaFold2 model of the GC domain of *Pf*GCα using the 2’-deoxy-3’-AMP of MamAC as an anchor point. These dockings suggest that the substrate and inhibitor adopt a mode of binding to *Pf*GCα comparable to that of 2’-deoxy-3’-AMP in MamAC, and the active sites appear relatively well conserved (Fig. 6D). Structural overlays indicate that the nucleobase occupies a highly similar position in the active sites (Fig. 6E). A few differences between the amino acids make it possible to accommodate a guanine base in *Pf*GCα instead of an adenine base in MamAC. Thus, while K938 in MamAC interacts with the N1 atom of adenine (as hydrogen bond acceptor), E3058 in *Pf*GCα can establish hydrogen bonds with the N1 atom and the amine function at the C2 position of guanine (as hydrogen bond donors) (Fig. 6D). MamAC D1018 interacts with the exocyclic amine of adenine, whereas the corresponding A3257 residue in *Pf*GCα allows no interaction with the oxygen in position C6 of guanine. Instead, this oxygen interacts with the main-chain amine of the adjacent residue (L3258). Baker (44) already highlighted that specificity for the nucleobase was due to the substitution of K578 and S684 in *Pf*AC to E3058 and A3257 in *Pf*GCα, respectively. We find that a third substitution seems necessary for guanine binding, as I940 of MamAC would clash with the exocyclic amine function of guanine (Fig. 6D). The corresponding residue is an isoleucine or a phenylalanine in other adenylate cyclases (I580 in *Pf*ACα). In *Pf*GCα the side-chain of this residue (V3060) is shorter, allowing the exocyclic amine of guanine to be accommodated. Finally, the two aspartates enabling interaction with phosphate groups (D396 and Asp440 in MamAC) are conserved in *Pf*GCα (D3975 and D4019). Interactions between *Pf*GCα and UA2239 are therefore predicted as plausible and show that UA2239 can replace GTP in the active site.

## Discussion

UA2239 belongs to the chemical family of ANPs. The advantages of these molecules lie in their chemical properties that make them very good candidates for the development of antimalarial agents. They are metabolically stable nucleotide analogs due to the presence of a phosphonate group, they can be synthesized in few steps from readily available and inexpensive chemicals, and they can be administered orally (*46, 47*). Surprisingly, the mode of action of UA2239 is totally different from those previously described for ANPs. When used in antiviral and anticancer therapies (*10, 34, 48*), the mechanism of action is linked to the inhibition of viral or cellular polymerases. In *Plasmodium*, already reported ANPs have been shown to inhibit the activity of recombinant enzymes of the purine salvage pathway (*31–33*). On the contrary, the extensive phenotypic characterization carried out in this study demonstrated that UA2239 targets the egress of merozoites from the iRBC and also inhibits gametocyte egress and male gametogenesis. The egress is a finely-tuned process under the control of *Pf*PKG activated by cGMP, whose level is regulated mainly by its synthesis *via Pf*GCα and its degradation *via Pf*PDEs. The observed decrease in cGMP in the presence of UA2239 provides evidence that *Pf*GCα is likely to be the primary target. Indeed, *Pf*GCα has been shown to be the only enzyme producing cGMP in asexual blood stages (*17*). This enzyme has also been shown to be essential for parasite growth and the effect of UA2239 on merozoite egress was similar to that observed when *Pf*GCα was disrupted (*17*). UA2239 is a GMP analog and our docking model shows that the compound can be accommodated and stabilized in the active site of the GC domain, supporting the idea that *Pf*GCα is the target of UA2239. Although the compound’s high selectivity for parasites is likely due to multiple factors, such as its ability to enter iRBCs and potentially accumulate, important amino acid differences between the *P. falciparum* and the mammalian enzymes may enhance its specificity for the parasite enzyme. In the catalytic sites, L3065 and A3257 of *Pf*GCα are not conserved in mammalian GCα but are both cysteine residues (*49*).

Despite this body of evidences, it cannot be ruled out that the drug may act on other regulators upstream of GCα activity. For example, the role of the adjacent P4-ATPase domain (*17*) or CDC50s (*50*) in regulating GC activity has not yet been fully elucidated. Interacting partners of *Pf*GCα have been identified like the unique GC organizer (UGO), and the signaling linking factor (SLF), and both play an important role in up-regulating GC activity just before egress in the asexual blood stage (*51*). In addition, GCα has been identified as a substrate of the protein phosphatase PP1 (*52*) and is most likely phospho-regulated (*53*). Clearly, further studies are needed to decipher, at the molecular level, the mechanism of action of UA2239 on its target.

Interestingly, resistance is not conferred by mutations in *Pf*GCα itself but in downstream effectors of the egress pathway. The effect of UA2239 on reducing cGMP levels can be circumvented by two resistance mechanisms. Indeed, mutations were found in the other two key enzymes of the cGMP signaling pathway, *Pf*PKG and *Pf*PDEβ. Mutations in *Pf*PKG were then validated as being responsible for the resistance mechanism. However, the mutated lines have an important fitness cost and are rapidly outcompeted by the wild-type strain in culture. While extrapolation from *in vitro* conditions to the field should be done with precaution it might be reasonable to assume that these mutations will not easily appear in the field. It is worth noting that the mutations in *Pfpkg* conferring resistance to UA2239 were not found in the 16,203 *P. falciparum* field isolates from 33 countries available in the *Pf*7 dataset using the *Pf*-HaploAtlas app (*54*). In addition, the minimum inoculum for resistance (MIR) for UA2239 was found to be of 3.4 × 10^8^ parasites (see methods section). These findings support the hypothesis that it can be difficult to generate resistance in the field.

If *Pf*PKG is not directly inhibited by UA2239, how can we explain the mechanism of resistance? *Pf*PKG is in an inactive conformation until the cytosolic level of cGMP increases. Cooperative binding of cGMP first to CNB-D, then to CNB-A and CNB-B changes the protein into its active conformation that *in fine* triggers the egress. This regulatory process is essential to ensure that merozoites egress neither too early nor too late in the cycle. Taking advantage of the available *Pf*PKG 3D structure (*39*), we analyzed the impact of the mutations R420I, H514N/Y and D597Y that all render the parasite resistant to UA2239. Interestingly, the three amino acids are all located at the interface between the CNB-D domain and the catalytic domain (Fig. 7). D597 of the catalytic domain and H524 of the CNB-D domain are both in interaction with amino acids of the opposite domain (*39, 55*). R420 is involved in water-mediated contacts with the C-terminus tail of the catalytic domain (*39*) and possibly in salt bridges with D597. All these interactions seem to be crucial to maintain *Pf*PKG in its inactive form in the absence of cGMP. Mutations of R420, H524 or D597 lead to a loss of these contacts and are expected to destabilize interactions between the CNB-D and the catalytic domain, thereby destabilizing the inactive form of *Pf*PKG. Modifications of these amino acids have already been studied. Franz *et al*. showed that a D597N mutation resulted in 3-fold decrease in the activation constant of *Pf*PKG, indicating that mutating this amino acid causes destabilization of the inactive state of the enzyme (*55*). An H524A mutation also destabilizes the inactive *Pf*PKG apo-form (*39*). We suggest that the mutations identified here destabilize *Pf*PKG that therefore requires less cGMP than wild-type *Pf*PKG to switch from the inactive apo-form to the active holo-form. The four mutations in *Pf*PKG are thus expected to confer resistance through a compensatory process.

**Fig. 7.**
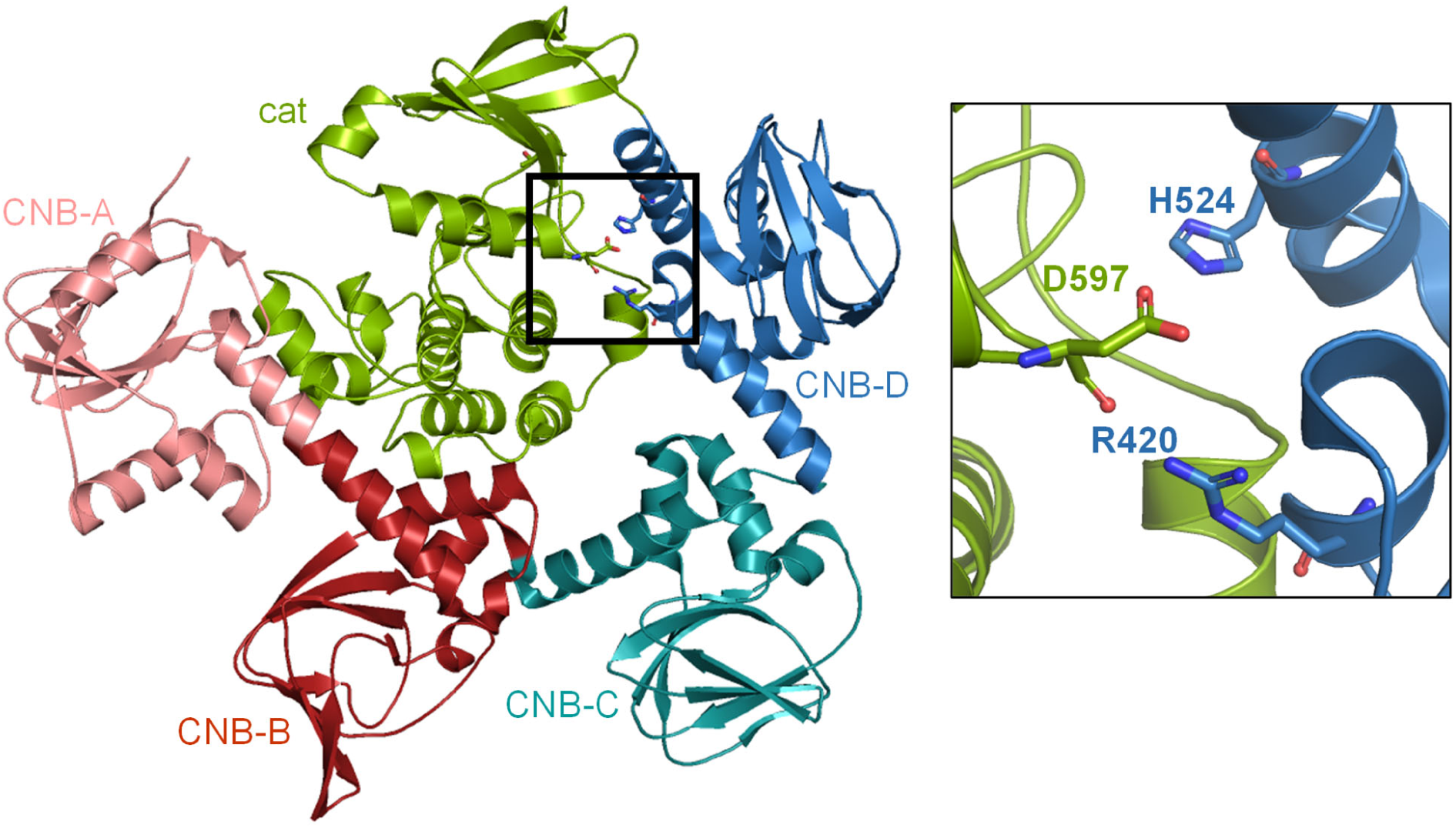
Localization of mutated amino acids on the 3D structure of *Pf*PKG. Left: X-ray 3D structure of the full length *Pf*PKG (PDB: 5DYK)^38^ showing the three mutation sites conferring resistance to UA2239 (black square) at the interface between the catalytic domain (green) and the CNB-D domain (blue). Right: close-up view of the interface showing the 3 mutated amino acids in the resistant strains. The colors of the domains are the same as shown in the schematic representation of the *Pf*PKG in Figure 4A.

The only resistant clone that did not have a mutation in the *Pfpkg* gene was mutated in the *Pfpdeβ* gene. Remarkably, the Y539D resistance mutation in *Pf*PDEβ is not localized in or close to the catalytic site but at the beginning of the fourth (of six) predicted transmembrane helices as shown in the AlphaFold model of the protein (fig. S8). At this stage we can only speculate that the mutation might induce a conformational change of PDEβ and/or prevent binding of potential partners, leading in one way or another to a decrease in enzymatic activity, resulting in higher levels of cGMP, which would then be sufficient to timely activate PKG in the presence of UA2239.

Decades of studies have demonstrated the importance of the cGMP-PKG dependent egress pathway in the malaria parasite and the potential for the involved proteins to be valuable therapeutic targets. Several PKG and PDE inhibitors have been extensively studied in recent years (*19–22, 26*), but to our knowledge UA2239 is the first GCα inhibitor. By disrupting the balance of cGMP in the parasite, which plays a crucial role at multiple stages of its life cycle, UA2239 interferes with the signaling pathway associated with this second messenger, leading to lethal effects. The pharmacological profile of UA2239 fits the criteria for promising antimalaria compounds as defined by Medicines for Malaria Venture (MMV). UA2239 is effective at low nM concentration against sensitive and chemo-resistant strains, is active *in vivo* in a mouse model and targets at least two stages of the parasite life cycle. Its effect on reducing the level of cGMP and its irreversible action on merozoite egress and on gametogenesis have been established. Its mechanism of action is new, although it has not yet been fully elucidated at the molecular level. UA2239 clearly provides a therapeutic advantage by irreversibly targeting parasite development and transmission to the mosquito, even being administered during the first days of gametocytogenesis. This study revealed the antimalarial potential of UA2239 and its novel mechanism of action, laying the foundation for developing an oral formulation with the aim of advancing it as a clinical candidate in the pipeline of antimalarial drugs.

## Materials and Methods

### Parasite culture

*P. falciparum* asexual blood stage parasites were grown in O^+^ or A^+^ erythrocytes supplied by the local blood bank (Etablissement Français du Sang, Pyrénées Méditerranée, France). Cultures were at 5% hematocrit in complete medium, constituted of RPMI 1640 containing 25 mM HEPES (Gibco Life Technologies), supplemented with 10% of human AB+ serum and 25 µg/ml gentamicin (Sigma). Cultures were maintained at 37°C under a controlled tri-gas atmosphere of 5% O_2_, 5% CO_2_ and 90% N_2_.

### *Plasmodium* growth inhibition assays

Drug effects on *P. falciparum* growth were measured in microtiter plates according to a modified Desjardins test (*9, 56*). *P. falciparum* infected erythrocyte suspension (1.5% final hematocrit and 0.6% parasitemia) were grown in complete medium with or without drug. The compounds were dissolved in RPMI 1640 or DMSO and then further diluted in culture medium. Parasite growth was assessed by measuring the incorporation of [^3^H]-hypoxanthine into nucleic acids as previously described (*57*). Suspensions of *P. falciparum*-infected erythrocytes were incubated with various concentrations of drug during 48 h. Then 30 µl of [^3^H]-hypoxanthine (0.5 µCi/well) were added for an additional 24 h-period. The reactions were stopped by freezing at -80°C. Cells were lysed by thawing and the parasite macromolecules including nucleic acids were recovered by harvesting the lysate on glass-fiber filter plates (Unifilter 96 GF/C, Perkin Elmer) using a FilterMate cell harvester (Packard Instruments). The radioactivity was counted on a Microbeta2 counter (Packard Instruments Revvity). Radioactivity background was obtained from incubation of non-infected erythrocytes under the same conditions and the value obtained was subtracted. Parasitemia were evaluated and expressed as percentage of the control (without drug). Results were expressed as IC_50_, which is the drug concentration leading to 50% inhibition of parasite growth and were determined by nonlinear regression analysis of the dose-inhibition curve using GraphPad Prism 8.3.0 software. Data were normalized to the level of incorporation in the untreated, and in the RBC (negative control). The experiments were performed at least three times, each at least in duplicates.

For stage-dependent susceptibility, drug was added at various concentrations to synchronized cultures at ring (0-3 hpi), trophozoite (18-21 hpi) or schizont (36-39 hpi) stages. After incubation for 6 h, cells were washed 3 times and resuspended in fresh complete medium without drug and incubated until 52 hpi. Then, [^3^H]-hypoxanthine (0.5 µCi per well) was added for 24 h and the reaction was stopped by freezing the plates at - 80°C. Cell viability was assessed following the above procedure for the drug sensitivity assay.

For time course assays, 750 nM UA2239 was added to synchronized cultures at the trophozoite (24 hpi) or schizont (36 hpi) stages. After incubation for various periods (30 min, 1 h, 2 h, 3 h, 4 h and 6 h), cells were washed twice and resuspended in fresh complete medium. [^3^H]-hypoxanthine was added at 52 hpi and incubated for an additional 24 h. Cell viability was assessed as described above.

### Interaction between drugs - Isobolograms

The interaction between UA2239 and other drugs, furosemide, chloroquine (CQ) and dihydroartemisinin (DHA), was studied using the isobologram method (*58*). The antimalarial activity of UA2239 was determined in the presence of the second drug at several concentrations that were lower than its IC_50_. The IC_50_ of each drug alone was found to be 100 µM, 11 nM and 2 nM for furosemide, CQ and DHA, respectively. In these experiments, compound concentrations were expressed as a fraction of the IC_50_ of the compounds alone. These fractions were called fractional 50% inhibitory concentrations (FIC_50_). The type of interaction was determined by calculating the maximal sum of FIC_50_ of the two compounds combined for antimalarial characterization (ΣFIC_50_ = FIC_50_ UA2239 + FIC_50_ drug B). Isobolograms were constructed from the FIC_50_s of UA2239 and drug B at the tested fixed concentration ratios. A straight line represents additivity (ΣFIC_50_ ≥ 1 and <2), a concave line denotes a trend toward synergy (ΣFIC_50_ < 1) or high synergy (ΣFIC_50_ ≤ 0.5), and a convex curve represents antagonism (ΣFIC_50_ ≥ 2) (*58, 59*). In all experiments, UA2239 was diluted in serial 1/3 dilutions while the second compound was added at several fixed concentrations.

### Follow-up of parasite morphology and DNA content

After tight synchronization by percoll-sorbitol method, 0-2 hpi rings (3D7) were treated with 2 µM UA2239. At intervals of 6 h over a period of 72 h, thin blood smears were performed and aliquots of infected erythrocytes were fixed in 4% paraformaldehyde (PFA) for 4 h at room temperature (RT) before storage at 4°C. Just before analysis, cells were washed twice with phosphate-buffered saline (PBS) and stained with 3.3⊆ SYBR Green I (Invitrogen) for 30 min. Cells were washed once and resuspended in PBS. Fluorescence was measured with a Becton Dickinson FACS Canto 1 cytometer and 100,000 events were recorded per sample. Data were analyzed with Diva software.

Cellular DNA content correlates with the intensity of fluorescence.

### Immunofluorescence assays

PV1-GFP (*35*) infected erythrocytes at late schizont stage were incubated with UA2239 (750 nM), C2 (1.5 µM) or E64 (10 µM) for 4 h. Then, thin blood smears were realized for each condition and cells on the smears were fixed with 4% PFA in PBS for 30 min at RT or overnight at 4°C in a humidified chamber, followed by 4 min of quenching with 0.1 M of glycine/PBS. Cells were then permeabilized with 0.1% Triton/PBS for 5 min and incubated for 30 min with 1.5% of bovine serum albumin (BSA)/PBS solution to block unspecific binding. Primary anti-GFP rabbit antibody (Invitrogen, A-6455) was added at 1:4,000 dilution for 1 h, then cells were washed 3 times with PBS before addition of

Alexa-488 goat antirabbit secondary antibodies (1:10,000; Invitrogen) for 1 h. Nuclei were stained with Hoechst (1:10,000). Slides were mounted with vectashield® medium for imaging. Images were taken on a Leica Thunder microscope at the Montpellier MRI imaging facility. Images were processed by Fiji software for sectioning and contrast adjustment.

### Electron microscopy

Highly synchronized *P. falciparum* cultures (0-3 h) were treated with 2 µM UA2239 at 6 hpi or 36 hpi. At 44-46 hpi, iRBCs were purified on VarioMACS columns (Miltenyi Biotec). As controls the same cultures were treated with C2 at 36 hpi or left untreated.

Samples were fixed by adding 25% glutaraldehyde (EM grade) directly in the culture medium in order to obtain a final concentration of 2.5%. After 10 min at RT, cells were centrifuged and the pellet resuspended in 20 pellet volumes of cacodylate buffer 0.1 M containing 2.5% glutaraldehyde and 5 mM CaCl_2_. The suspension was left 2 h at RT and then kept at 4°C in the fixative until further processing. All the following incubation steps were performed in suspension, followed by centrifugation. Cells were washed with cacodylate buffer and post-fixed with 1% OsO_4_ and 1.5% potassium ferricyanide in cacodylate buffer. After washing with distilled water, samples were incubated overnight in 2% uranyl acetate in water and dehydrated in graded series of acetonitrile. Impregnation in Epon 812 was performed in suspension in Epon:acetonitrile (50:50) and two times 1 h in 100% Epon. After the last step, cells were pelleted in fresh Epon and polymerized 48 h at 60°C. All incubation and washing steps were performed using a microwave processor PELCO Biowave Pro+ (TED PELLA) except for the overnight incubation. 100 nm sections were made with an ultramicrotome Leica UCT collected on silicon wafers, contrasted with uranyl acetate and lead citrate and observed on a Zeiss Gemini 360 scanning electron microscope on the MRI EM4Bio platform under high vacuum at 1.5 kV. Final images were acquired using the Sense BSD detector (Zeiss) at a working distance between 3.5 and 4 mm. Independent random tiles were acquired with a pixel size of 5 nm and red blood cells infected with segmented schizonts were counted. Representative pictures were acquired with a pixel size of 1 nm for illustration.

### Effect on *in vitro P. falciparum* gametocytogenesis and gametogenesis

*P. falciparum* parasites of the B10 clone (*60*) were cultivated under standard conditions using RPMI 1640 medium supplemented with 10% heat-inactivated human serum and human erythrocytes at a 5% hematocrit. Parasites were kept at 37°C in a trigas atmosphere. Synchronous production of gametocytes was achieved by treating synchronized cultures at the ring stage (10-15% parasitemia, day 0) with 50 mM *N*-acetyl glucosamine (NAG) to eliminate asexual parasites. NAG was maintained for 5 days until no asexual parasites were detected in the culture. At day 1 post NAG addition, 750 nM UA2239 were added to the medium either for 24 h or for 10 days. Giemsa-stained smears were scored daily for gametocyte density and distribution of gametocyte stages (stages I to V). At day 10 post NAG, stage V gametocytes were subjected to gametogenesis assay (*8*). In brief, 200 µl samples from each condition were stained for 15 min at 37°C with 5 µg/ml Wheat Germ Agglutinin (WGA)-Alexa Fluor 647 and 0.5 µg/ml DNA intercalating dye Hoechst 33342. Samples were then pelleted at 1,000 × *g* for 1 min, then mixed in 50 µl human serum and subjected to a temperature drop (from 37°C to 22°C/ RT) for 20 min during which gametogenesis took place. Samples were fixed either before or 20 min after activation of gametogenesis with 1% PFA for 30 min at RT. After one wash in PBS, stained cells were mounted in a microscope slide under a sealed coverslip. Samples were observed at 100× magnification using a Leica DMi8. At least 100 gametocytes were analyzed for each sample in three independent experiments. Percent rounding and gamete egress were determined by calculating the percentage of round gametes and of WGA-negative gametes (due to loss of host erythrocyte) in the total gametocyte population, respectively.

### Effect on gametogenesis after treatment of *P. berghei* infected mice

Gametogenesis was evaluated by counting the number of exflagellation centers in blood of female NMRI mice infected with *P. berghei* ANKA at a parasitemia of 15 - 30%. For each mouse, a drop of blood was taken from the tip of the tail and placed between a slide and a coverslip, before (control) and 30 min after intraperitoneal treatment with UA2239. After 10 min at 20°C, exflagellation centers were counted under the microscope and the number of exflagellation centers per 10,000 cells was determined. The counting was done twice for the control before UA2239 treatment. Activity of UA2239 is expressed as the number of exflagellation centers/10,000 cells as a percentage of control.

### Generation of UA2239 resistant strains and analysis of whole genome sequencing

We used two different protocols to obtain UA2239-resistant *P. falciparum* blood-stage parasites. Drug pressure was applied intermittently by drug-on/drug-off cycles either using the same high concentration (10⊆IC_50_) or gradually increasing the concentration of UA2239 (from 3⊆ to 10⊆IC_50_). Parasites were cultured in presence of UA2239 until parasites were no longer detectable on thin blood smears. The cells were then washed and cultured in the absence of drug until parasitemia reached 1% and another cycle of drug pressure began. After each cycle, parasite sensitivity to UA2239 was determined. Parasites were cultured independently and in triplicates for 7 months. From all six independent and resistant populations, several clones were isolated. The 3D7 wild-type strain was cultured in parallel during the 7 months and was also included in the whole-genome sequencing, but as a population. Sequencing was done by MGX-Montpellier GenomiX in two independent runs on Illumina MiniSeq platform. The 150 bp paired-end sequencing reads from both runs were merged for each sample to increase the sequencing depth. Sequence adapters were trimmed with Trimmomatic (version 0.39) and filtered reads were mapped on the 3D7 reference genome (version 46) with bwa 0.17 (*61, 62*).

Single-nucleotide polymorphisms (SNPs) and micro insertions/deletions (INDELs) were discovered with GATK (version 4.2.0.0) HaplotypeCaller, CombineGVCFs and GenotypeGVCFs (*63*). SNPs and micro INDELs were discarded from a sample if they were covered by less than 5 reads. As each sequenced sample originates from a clonal population, only homozygous variant calls (at least 90% of the sample reads supporting one allele) were considered. SNPs and micro INDELs with a mutated allele present in a drug-resistant sample and absent from the drug-sensitive sample were extracted using varif (https://github.com/marcguery/varif, version 0.5.4). Macro INDELs, duplications, inversions and translocations were discovered and filtered with DELLY (version 0.8.7) using the options ‘-p -f somatic -a 0.5 -r 0.5 -v 10 -c 0.01’(*64*). Haplotypes of selected proteins of interest were screened with *Pf*-HaploAtlas in the 16203 *Plasmodium falciparum* samples available in the Pf7 dataset (*54, 65*).

### Generation by CRISPR-Cas9 of transgenic 3D7 lines with single amino acid changes in *Pf*PKG

The *Pf*3D7 *pkg* locus consists of 5 exons spanning 3561 base pairs (bp) encoding a protein of 853 amino acids. The mutations R420I, H524N/Y, D597Y identified in the UA2239-resistant strains by the WGS and a previously identified T618Q mutation rendering parasites resistant to the PKG inhibitor C2 are all located in exon 5 (fig. S3). A sequence of 675 bp covering all these positions (position 2212 to 2886 of the *pkg* locus) was recodonised (termed ePKG). Six synthetic DNA fragments were ordered, five of which encode a single of the above listed mutations and one without mutation. Homology regions HR1 (510 bp) and HR2 (658 bp) for double homologous recombination were both cloned together by In-Fusion cloning (Takara-Bio) in a KpnI/SacI digested pBluescript plasmid generating an EcoRV site between HR1 and HR2 that was subsequently used to insert in frame by In-Fusion cloning the recodonised sequences. All plasmids were verified by Sanger sequencing.

Three different guide RNA sequences (g12, g13, g15) were identified in the 675 bp sequence using ChopChop (*66*). Complementary primer pairs were designed, annealed and ligated into BbsI digested plasmid pDC2-cas9-sgRNA-hDHFR/yfcu (*67*). The pDC2 plasmid and the pBluescript construct (75 µg each) were transfected together using ring stage cultures of 3D7 at about 5% parasitemia using a BioRad Gene Pulser (950 V, 310 µF, 200 Ω) (*68*). Selection with 2.5 nM WR22910 (a gift from Jacobus Pharmaceuticals) was started 5 h after transfection and was maintained during daily media changes for 1 week. Genome edited parasites were obtained with one of the three guide RNAs used.

Parasite populations were cloned by limiting dilution.

### Growth competition assay

To quantify the growth of GER-parasites in competition with 3D7 wild-type parasites, each GER strain was mixed with the 3D7 wild-type at equal parasitemia in 5 ml dishes and maintained in culture for a total of 40 days. Aliquots were collected regularly and DNA was extracted using the DNA blood extraction kit (Qiagen). qPCR was done using the LightCycler 480 Sybr Green I system (Roche) and the LightCycler 480 Software, Version 1.5 was used for “absolute fit point” data analysis. The strains could be distinguished through the presence of recodonised *epkg* sequences in all GER parasite lines. Forward primer AGGTGCCACGGCCGATTATGC and reverse primer GCTACTGTATGATGAACCGC were specific for all GER-parasites. Forward primer CAAGGACCTATGTTAGCACATTTG and reverse primer CTAATTTAACTGTTCCGAAAGTACCTCTT were specific for 3D7 *Pfpkg* sequence. To validate that the presence of recodonised sequence in the *pkg* gene did not in itself induce a fitness cost, we also performed a competition assay between 3D7 and the ePKG-WT GER parasite clone. No effect of recodonisation on the parasite fitness was observed (fig. S9).

### Ring-stage survival assay (RSA)

After tight synchronization by percoll-sorbitol method, 0-3 hpi rings were diluted to 0.5 - 1% parasitemia and 2% hematocrit as previously described (*69*). Parasites were exposed to 0.01% of DMSO (control) or to 700 nM DHA for 6 h at 37°C in a tri-gas atmosphere in duplicates for each condition and strain. Then the cultures were washed and cultivated for additional 66 h. Culture medium was changed daily. The parasitemia was determined on blood smears by counting at least 10,000 cells. Survival rates were calculated as the ratio of parasites in treated and untreated cultures. A strain is considered resistant to artemisinin if the survival rate is higher than 1%. As positive control for resistance the *P. falciparum* NF54 strain with the C580Y substitution in Kelch 13 (*70*) was used (gift from Jose-Juan Lopez-Rubio). *P. falciparum* ART-sensitive 3D7 and NF54 strains were used as negative controls.

### Enzymatic assay of recombinant *Pf*PKG

The full-length *Pf*PKG was expressed and purified by the Protein Biochemistry Platform of the faculty of medicine, Geneva University. Baculoviruses expressing the *Pf*PKG were generated from a modified pFastBac vector like vector for insect cell expression. It contains an N-terminal “TwinSTREP-His-TEV” tag. Protein samples were first purified by affinity using STREP-tactin resin. A second step of purification was done after TEV cleavage by size exclusion chromatography using a Superdex 200 Gel Filtration column equilibrated in 20mM Tris pH 7.4, 150mM NaCl and 3mM DTT. *Pf*PKG inhibition assays were performed based on previously described methods (*21, 22*), using ADP formation as a measure of kinase activity. The reaction buffer for all measurements consists of 50 mM Tris-HCl pH 7.5, 40 mM KCl, 10 mM MgCl_2_, 5 mM β-mercapto-ethanol, 10 µM cGMP, 1 mM ATP, 50 µM peptide substrate GRTGRRNSI (GeneCust) and 100 nM purified *Pf*PKG protein. UA2239 was added or not at 10 µM or 100 µM and C2 at 10 µM. Controls were done in absence of enzyme or cGMP. Reactions were initiated by the addition of ATP and cGMP, and incubated for 45 min at RT. ADP formation was measured using the ADP-Glo Kinase Kit (Promega) according to manufacturer’s instructions. Briefly, 20 µl ADP-Glo Reagent was added to 20 µl kinase reaction in white 96 well plates (Perkin Elmer) and incubated for 40 min at RT to deplete the remaining ATP. Then, 40 µl of kinase detection reagent was added and the reaction was incubated for a further 60 min at RT. Luminescent signal was measured using the Sunrise Microplate Reader (Tecan). Reactions were carried out in triplicates.

### Measurement of intracellular cGMP levels

cGMP levels were measured using Direct cGMP ELISA kit (Enzo Life Sciences). Mature schizonts at 36 hpi were purified on VarioMACS columns. Then approximately 1.5 ⊆ 10^8^ parasites were incubated in presence of 1.5 µM C2 (to prevent schizont rupture), without or with various UA2239 concentrations. At 42 hpi, parasites were collected by centrifugation (1 min at 600⊆g) and resuspended in 100 µl of 0.1 M HCl. After 3 min of incubation at RT, samples were centrifuged for 5 min at 9,000⊆g. Supernatants were collected, and diluted by addition of 600 µl of 0.1 M HCl, and frozen at -80°C. cGMP levels were measured with the acetylated protocol according to the manufacturer’s instructions. Absorbance was detected at 405 nm on a Spark spectrophotometer (Tecan). The data were analyzed using GraphPad Prism 8.3.0. To validate experiments, the PDE inhibitor BIPPO was added to samples with C2 only or with C2 and 750 nM UA2239 at 42 hpi for 3 min and then processed as the other samples.

### Treatment of parasite cultures with cGMP analog to rescue UA2239 treatment phenotype

Synchronous late schizonts were treated with 750 nM UA2239 or 1.5 µM C2 from 38 hpi. At 44 hpi, 60 µM PET-cGMP were added or not to both cultures. Giemsa-stained thin blood smears were taken after 2 h, and parasitemia of segmented schizonts and rings were determined (at least 5,000 cells counted).

### Generation of the *Pf*GCα 3D model and docking of GTP and UA2239

The structure of the guanylate cyclase domain of *Pf*GCα (residues 2741 to 4226) (fig. S8) was modelled using AlphaFold2 implemented on the CBS local server. The cyclase domain of this model was aligned with that of MamAC (PDB 1CS4, chains A and B), as shown in figure S8B-D. The R.M.S.D. between the cyclase domains was calculated from residues 3007-3314 and 3963-4151 of *Pf*GCα guanylate cyclase domain, excluding two insertions. Dockings of UA2239 and GTP into the guanylate cyclase domain of *Pf*GCα were performed with PLANTS 1.2 (*71*), using the coordinate of the C1’ atom of 2’-deoxy-3’-AMP in the 1CS4 structure and a shape constraint of -3. The binding site radius was fixed to 10 Å. The ten best-ranked poses were similar, except for the positions of the phosphates and the ribose for GTP which fluctuates (not shown).

### Determination of minimum inoculum of resistance

UA2239 resistance selection was performed by culturing 3D7 parasites continuously under 3 ⊆ IC_90_ (600 nM) drug pressure at 3% hematocrit in Petri dishes as previously described (*72*). Starting parasite inocula were 1.28 ⊆ 10^7^ (3 dishes) or 1.28 ⊆ 10^8^ (5 dishes). Drug-containing media was replaced every day for the first 5 days, then every 3 or 4 days. RBCs were replenished by half once a week. Cultures were monitored by Giemsa staining and microscopy daily until parasites were cleared, then 2 times per week to detect recrudescence. Selections were maintained for 60 days or until recrudescent parasites were observed. The MIR value is defined as the minimum number of parasites used to obtain resistance and calculated as follows: total number of parasites inoculated divided by the total number of positive cultures. This formula includes lower inocula where there were no positive wells and excludes higher inocula in cases where lower inocula already yielded resistance. In this experiment, we observed recrudescence in only two dishes of 1.28 ⊆ 10^8^ parasites, consequently the MIR value was calculated as follows: *MIR* = 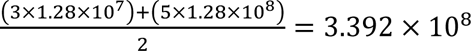 parasites.

### Statistical analysis

All statistical tests were performed using GraphPad Prism version 8.3.0. One-way ANOVA was performed for Fig. 3E. Two-tailed unpaired Student’s t-tests were performed for Fig. 3F, 6A, 6B and fig. S7. Paired-ratio Student’s t-tests were performed for Fig. 6C. For whole genome sequencing, all the samples were sequenced in two independent runs and both runs were merged for each sample to increase the average depth. For the effect of UA2239 on the rupture of the PVM by electron microscopy, 72 to 195 segmented schizonts were analyzed. For the effect of UA2239 on *P. falciparum* gametocytes development, at least 100 gametocytes were analyzed for each sample in three independent experiments.

## Acknowledgments

We are grateful to E. Jublanc, V. Diakou, and the imaging facility MRI at the University of Montpellier, as well as the cytometry facility MRI, both part of the national infrastructure France-BioImaging supported by the French National Research Agency (ANR-10-INBS-04, «Investments for the future»), for their assistance and technical support. We also thank P. Clair at the qPCR facility of University of Montpellier / Montpellier GenomiX for his technical advice. Whole genome sequencing was performed at the MGX-Montpellier GenomiX facility. The authors acknowledge the financial support from the France Génomique National Infrastructure, funded as part of “Investissement d’Avenir” program managed by the Agence Nationale de la Recherche (contract ANR-10-INBS-09). We thank M. Brochet, O. Vadas, L. Falconnet and R. Visentin from the Protein Biochemistry platform of the Geneva University for expression and purification of *Pf*PKG recombinant protein, J.J. Lopez-Rubio (LPHI, University of Montpellier) for providing Artemisin-resistant NF54 strain and O. Bilker (Umeå University) for C2 compound. We thank the ChemBioFrance research infrastructure and the French Infrastructure for Integrated Structural Biology (FRISBI) supported by the Agence Nationale de la Recherche (contract ANR-10-INBS-05).

## Author contributions

MA, RD, KW, SW designed, performed, and analyzed experiments on *in vitro* asexual *P. falciparum* stages and in *P. berghei*-infected mice. EC-I and CLa designed, performed, and analyzed experiments on *P. falciparum* gametocytes. M-AG and AC performed whole-genome sequencing data processing and analysis. LR and LB designed, performed and analyzed electron microscopy experiments. CLi has made the model of the GCα and the docking of GTP and UA2239. TC and SP synthesized UA2239. RC, SW, CLa., AC, and SP supervised the experiments. RC, SW conceived the study and wrote the manuscript. All authors approved the final version.

## Data and materials availability

All data needed to evaluate the conclusions in the paper are available in the main text and/or the supplementary materials. UA2239 is available after signature of a material transfer agreement.

