## Supplemental-data for "An Acyclic nucleoside phosphonate effectively blocks the egress of the malaria parasite by inhibiting the synthesis of cyclic GMP"

Marie Ali *et al.*

**This PDF file includes:**

Figs. S1 to S9
Table S1
Reference (#73)

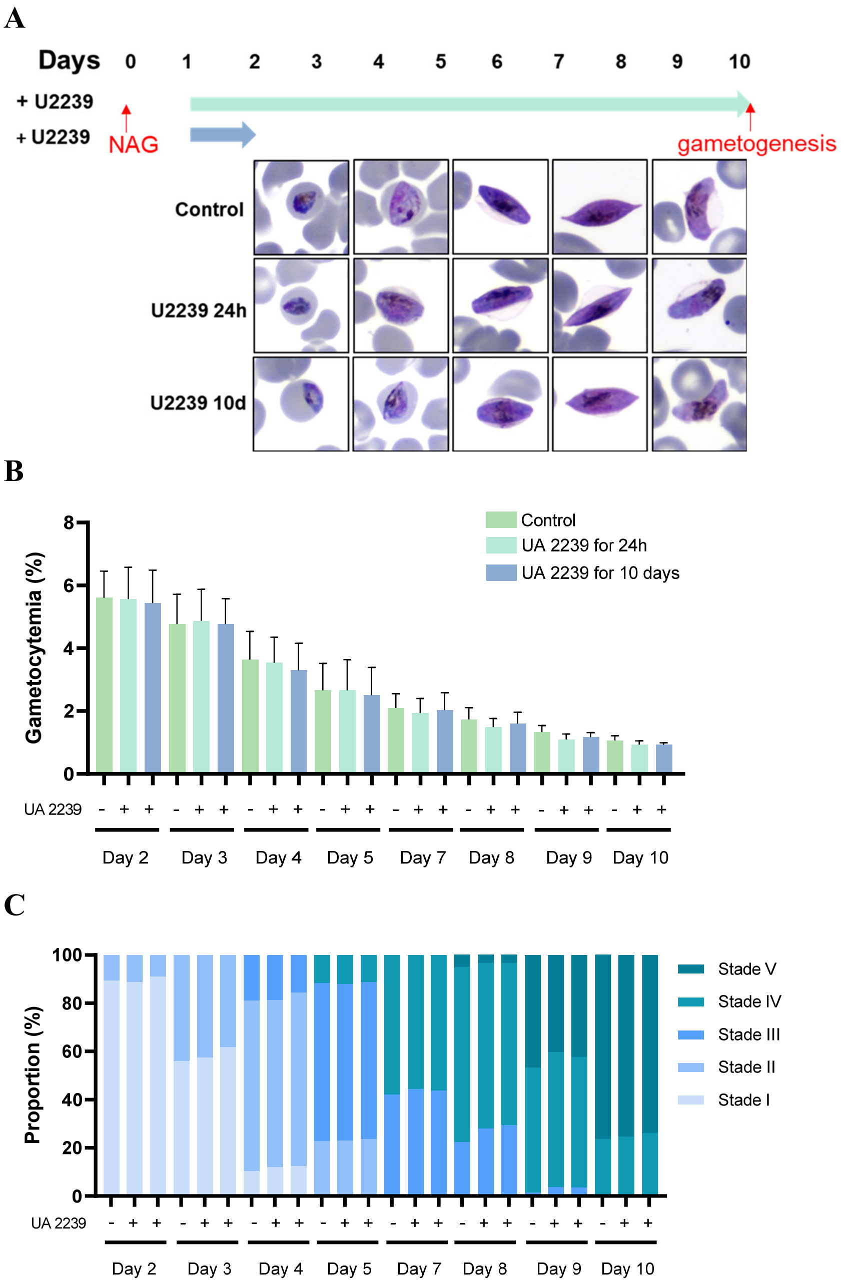


**Fig. S1. Effect of UA2239 on gametocytemia.** UA2239 was added or not on B10 cultures (NF54-derived clone) at day 1 post N-acetyl glucosamine (NAG) for 24 hours or for 10 days. Gametocytemia was followed during the 10 days for the 3 conditions. **(A)** Giemsa-stained smears of treated and untreated gametocytes. **(B)** Gametocytemia. **(C)** Proportions of the different stages of gametocytes up to 10 days of maturation (conditions in the same order as in B).

A

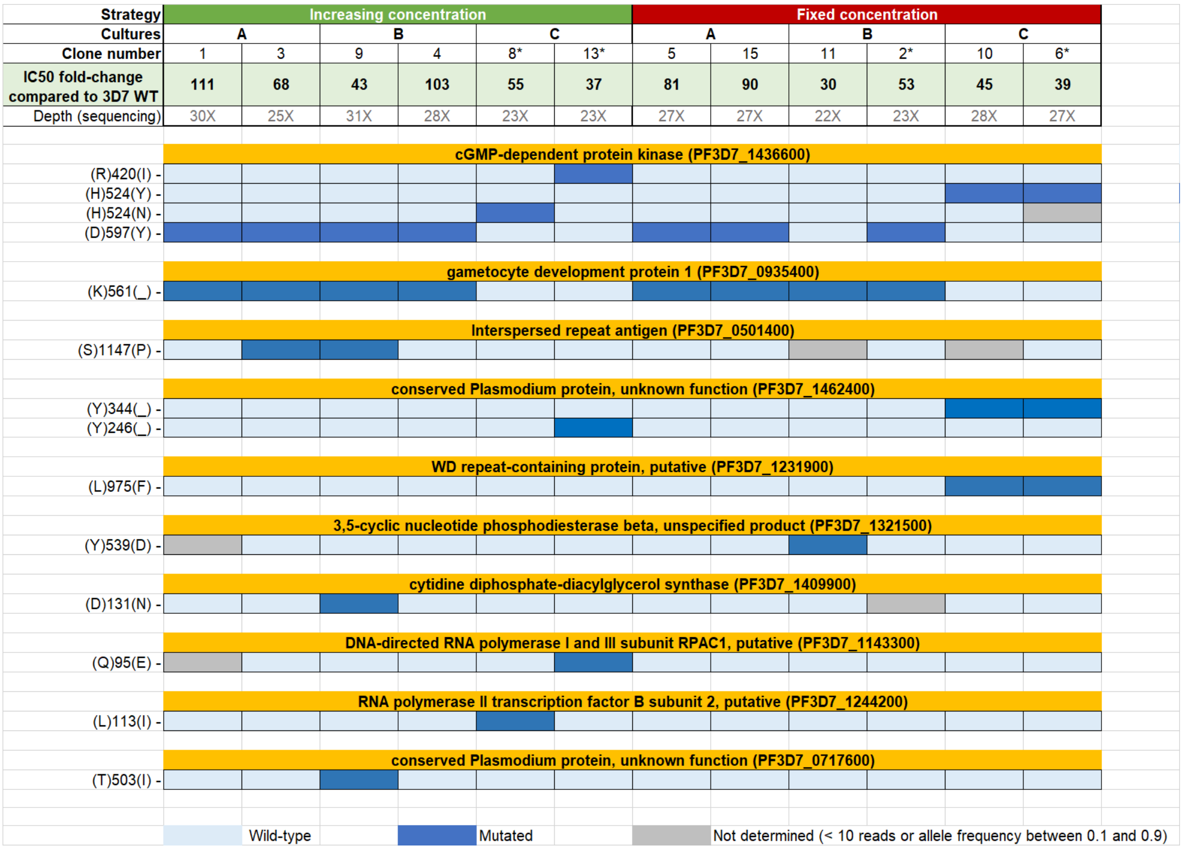
Strategy
Increasing concentration
Fixed concentration
Cultures
A
B
C
A
B
C
Clone number
1
3
9
4
8*
13*
5
15
11
2*
10
6*
IC50 fold-change compared to 3D7 WT
111
68
43
103
55
37
81
90
30
53
45
39
Depth (sequencing)
30X
25X
31X
28X
23X
23X
27X
27X
22X
23X
28X
27X
cGMP-dependent protein kinase (PF3D7_1436600)
(R)420(I) -
(H)524(Y) -
(H)524(N) -
(D)597(Y) -
gametocyte development protein 1 (PF3D7_0935400)
(K)561(_) -
Interspersed repeat antigen (PF3D7_0501400)
(S)1147(P) -
conserved Plasmodium protein, unknown function (PF3D7_1462400)
(Y)344(_) -
(Y)246(_) -
WD repeat-containing protein, putative (PF3D7_1231900)
(L)975(F) -
3,5-cyclic nucleotide phosphodiesterase beta, unspecified product (PF3D7_1321500)
(Y)539(D) -
cytidine diphosphate-diacylglycerol synthase (PF3D7_1409900)
(D)131(N) -
DNA-directed RNA polymerase I and III subunit RPAC1, putative (PF3D7_1143300)
(Q)95(E) -
RNA polymerase II transcription factor B subunit 2, putative (PF3D7_1244200)
(L)113(I) -
conserved Plasmodium protein, unknown function (PF3D7_0717600)
(T)503(I) -
Wild-type
Mutated
Not determined (< 10 reads or allele frequency between 0.1 and 0.9)

**B**

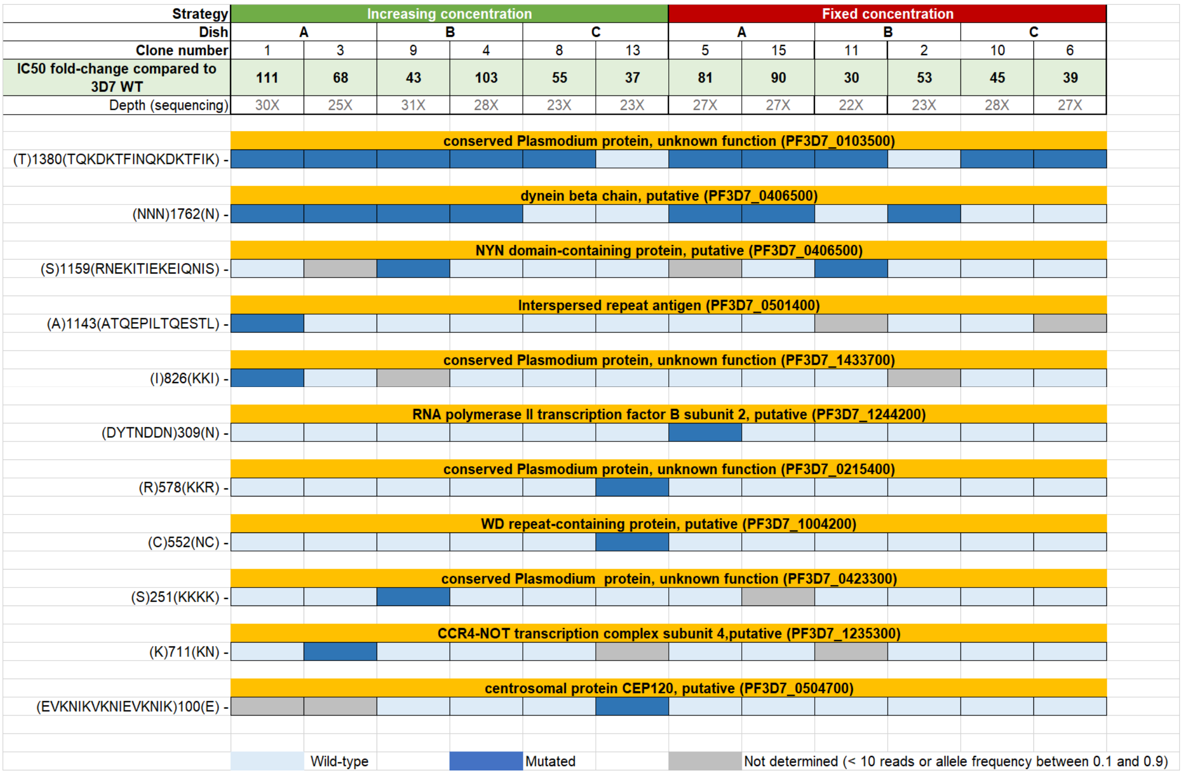


**Fig. S2. Selection of clones and whole genome sequencing of resistant parasites.**
Two clones per population of resistant parasites (R-parasites) were selected for whole-genome sequencing (12 clones in all), along with the 3D7 strain maintained in culture for the period required to obtain resistant parasites (sensitive parasites). Clone numbers are indicated, as well as the method used to obtain resistant parasites (exposure to an increasing or fixed concentration of drug), their $IC_{50}$ fold-change (ratio of $IC_{50}$ between resistant and WT parasites) and sequencing depth. Variants (SNPs and micro INDELs) detected in the 13 combined samples were reduced according to their presence in the drug-resistant clones and their absence in the drug-sensitive parasites. **(A)** 14 single nucleotide polymorphisms (SNPs) leading to non-synonymous mutation in one gene. The four clones mutated in the *Pfpkg* gene, chosen for further studies, are labelled with an asterisk. **(B)** 11 micro INDELs.

A

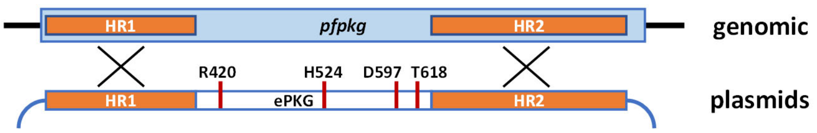


B

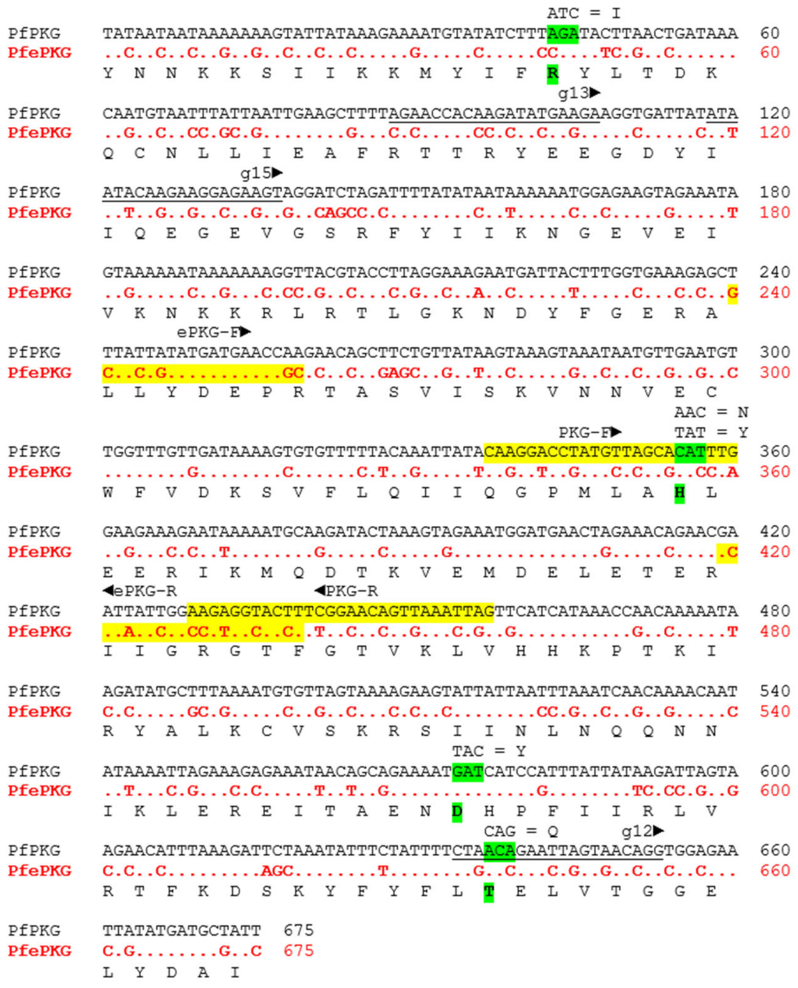

**Fig. S3. Generation of transgenic parasites carrying mutations in *Pf*PKG. (A)** Schematic representation of the last exon (exon 5, blue box) of the *Pfpkg* locus (above) and of the transfection constructs (below). The homology regions (HR) for double homologous recombination and the position of the mutated residues are indicated. The recodonised sequence is named ePKG. **(B)** Alignment of the *Pfpkg* wild type sequence (top) and the ePKG in red (below). Dots indicate identical bases. The amino acid sequence is shown at the bottom. Mutated residues are highlighted in green and the modified sequence in the mutants shown above. The three guide RNA sequences (g12, g13, and g15) that led to the generation of the mutants are underlined. Primer sequences that were used in PCR to differentiate between wild type and transgenic parasites are indicated in yellow.

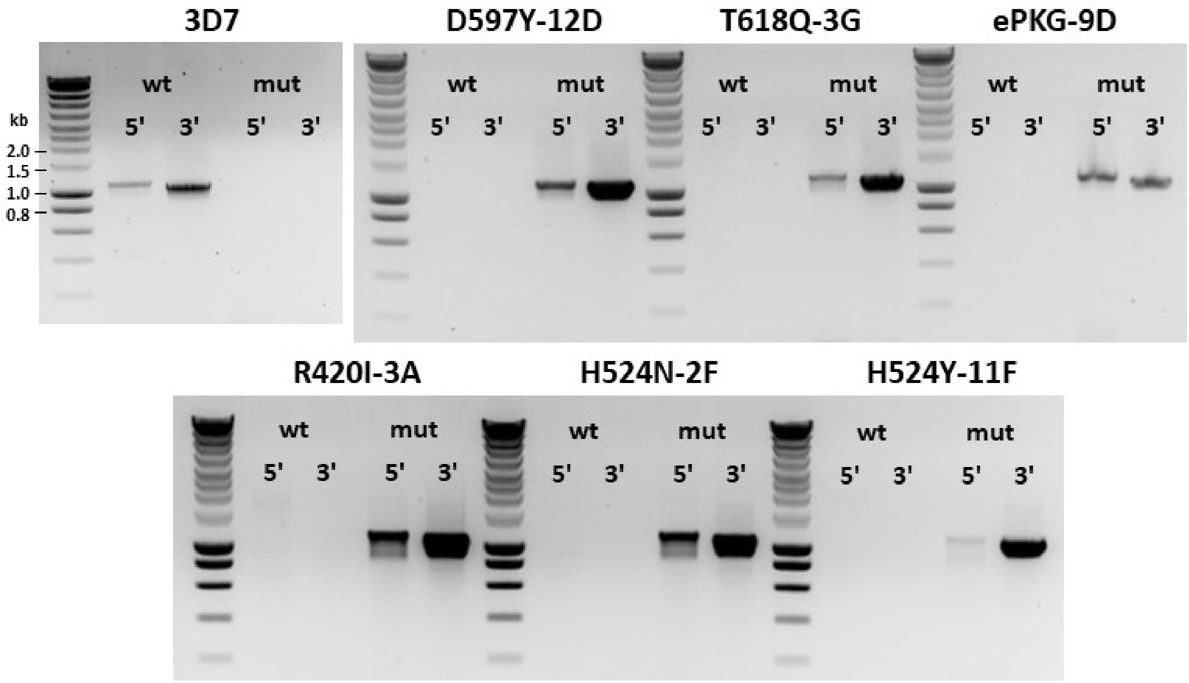


**Fig. S4. Genotyping of transgenic parasite clones carrying mutations in *Pf*PKG.** Correct 5'- and 3' homologous recombination was detected by PCR on genomic DNA. For 5' integration primer Int-F in the sequence upstream of the HR1 region was used in combination with either the PKG-R (wild type) or ePKG-R (mutant, recodonised) primer (see fig. S3). For 3' integration primer Int-R in the sequence downstream of HR2 was used in combination with either PKG-F (wt) or ePKG-F (mut). ePKG-9D corresponds to the transgenic clone carrying the recodonised sequence without any encoded amino acid change. SmartLadder (Eurogentec) was used as DNA molecular weight marker and relevant bands are indicated in kilobases (kb).

**A**

| Strain | IC$_{50}$ of CQ (nM) | |
|---|---|---|
| WT 3D7 | 11.4 ± 5.3 | |
| | **R-parasites** | **GER-parasites** |
| *Pf*PKG - R420I | 16.7 ± 7.2 | 12.0 ± 4.4 |
| *Pf*PKG - H524N | 17.5 ± 3.4 | 21.0 ± 8.7 |
| *Pf*PKG - H524Y | 12.3 ± 7.9 | 12.9 ± 3.2 |
| *Pf*PKG - D597Y | 12.2 ± 1.6 | 8.2 ± 3.9 |

**B**

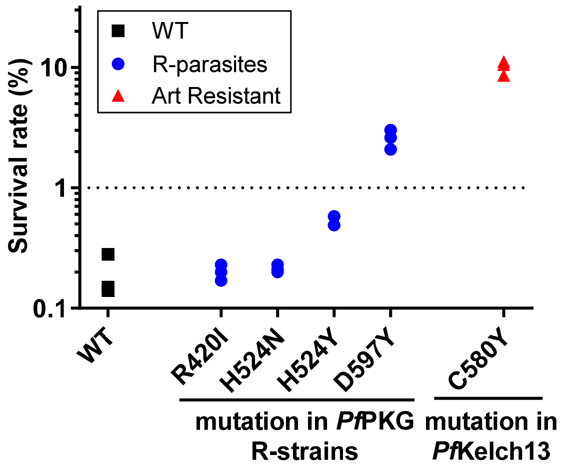


**Fig. S5. Sensitivity of resistant parasites to antimalarial compounds. (A)** Sensitivity of UA2239-resistant strains (R and GER-parasites) to chloroquine (CQ). Data are mean ± SD of 3 independent experiments performed in duplicates **(B)** Sensitivity of R-parasites to dihydroartemisinin (DHA). Sensitivity was determined using ring-stage survival assay. A survival rate <1% indicates sensitive strains. 3D7 WT served as the sensitive control and the NF54 Kelch13 C580Y as resistant control. Data are from 3 independent experiments.

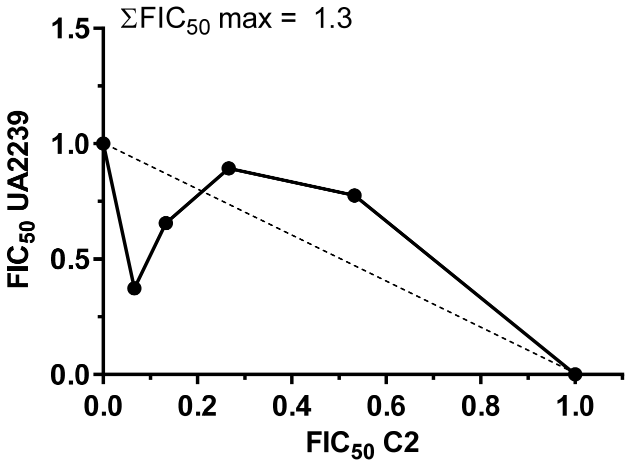


**Fig. S6. Interaction between UA2239 and C2.** *In vitro* interaction between UA2239 and C2 for their antimalarial activity is represented as an isobologram. The maximal sum of FICs (ΣFIC50 max) is indicated. Data presented are from one representative experiment (n=3 independent experiments).

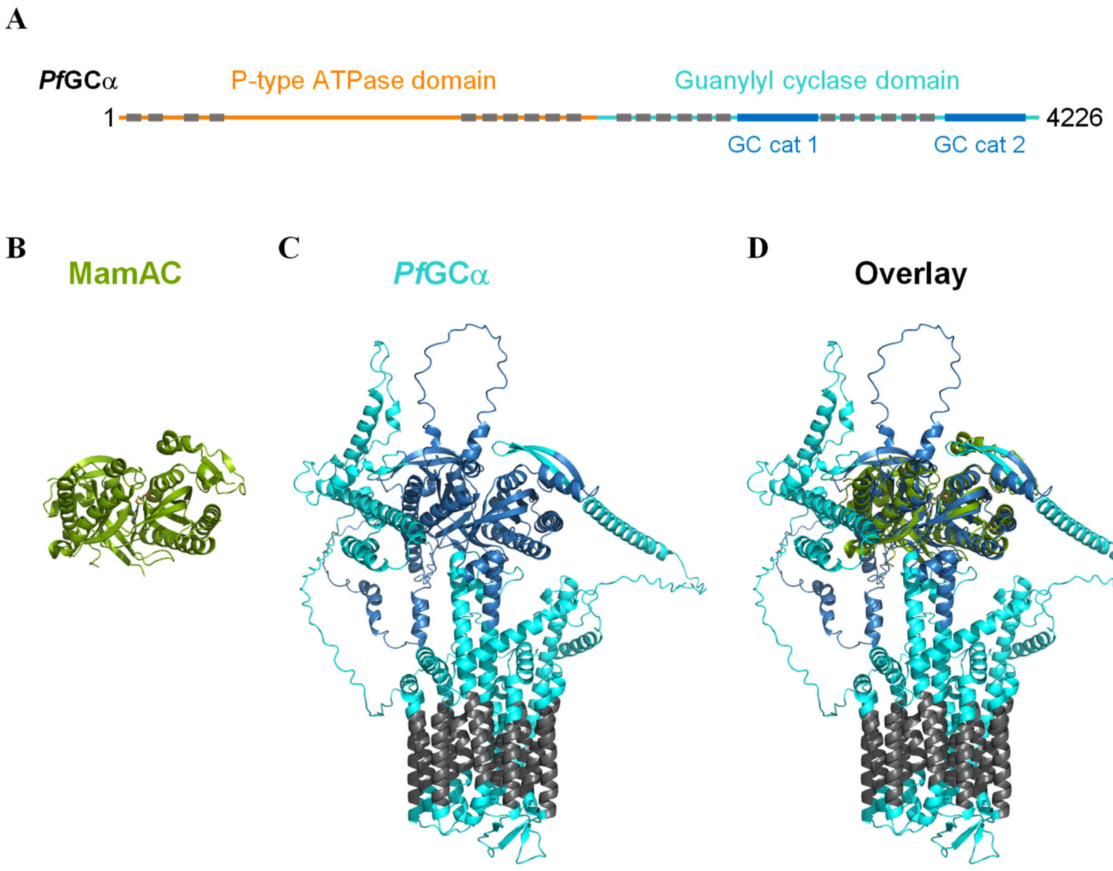


**Fig. S7. Model of the 3D structure of *Pf*GCα. (A)** Schematic representation of the full-length *Pf*GCα. The guanylate cyclase domain (residues 2741-4226) is shown in cyan and the two catalytic sub-domains (residues 3007-3314 and 3963-4151) are in dark blue. The position of the transmembrane helices as predicted by DeepTMHMM (*73*) are shown with grey boxes. **(B)** Structure of the adenylyl cyclase domain of MamAC determined by X-ray diffraction at 2.50 Å (PBD 1CS4). Chains A and B, forming the adenylyl cyclase domain, are shown in green cartoons. **(C)** AlphaFold model of the guanylate cyclase domain of *Pf*GCα shown in cartoons, using the same color code as in (A). **(D)** Overlay of MamAC and *Pf*GCα structures, aligned on the cyclase domains.

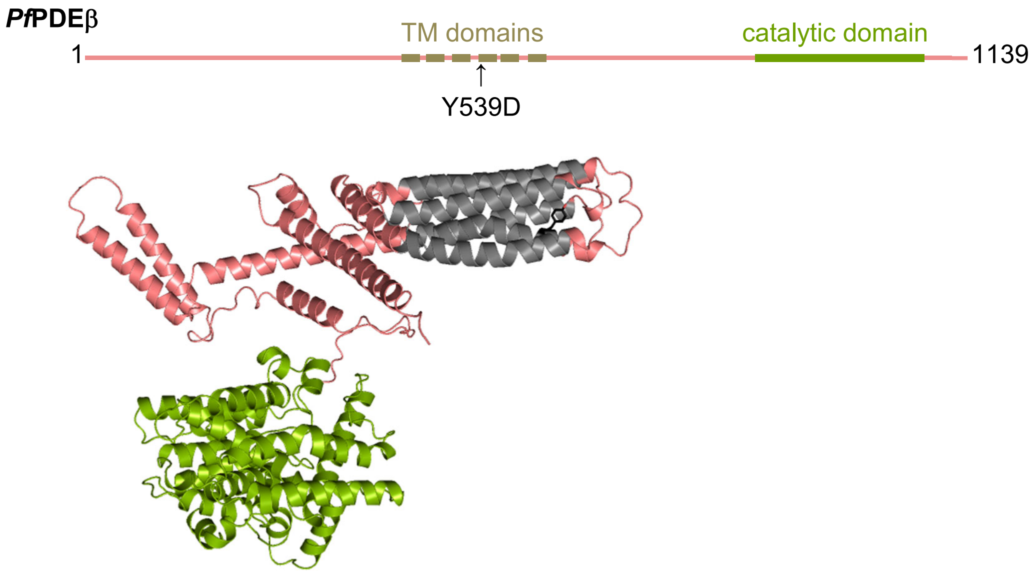


**Fig. S8. Schematic representation and AlphaFold model of *Pf*PDEβ** (PDB computed structure: AF-Q8I6Z7-F1-v4). The catalytic domain is shown in green and the 6 predicted transmembrane α-helices are in grey. The Y539 located in the 4th helix that is mutated in the resistant parasite line is in black. The structural model shows amino acid 342 to 1139.

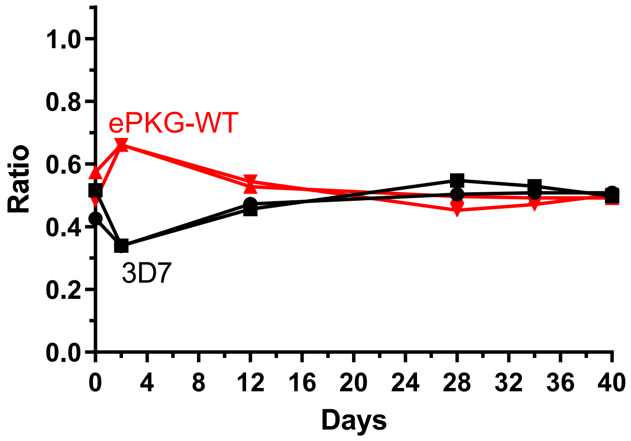


**Fig. S9. Pairwise competitive growth assay followed by qPCR between ePKG-WT and 3D7 WT strains.** Cultures for the *Pf*PKG competition assays were started at approximately equal parasitemia for both strains, equivalent to a ratio of around 0.5. The ratio remains at 0.5 if both strains grow at the same rate. The ratio rises to 1 if one strain grows faster than the other and finally represents 100% of the parasites (n=2).

| Treatment | Number of segmented schizonts analyzed | % of intact PVM |
|---|---|---|
| Untreated | 72 | 90.3% |
| UA2239 from 6 hpi | 116 | 88.8% |
| UA2239 from 36 hpi | 195 | 98.0% |
| C2 from 36 hpi | 139 | 82.0% |

**Table S1. Percentage of parasites whose PVM did not rupture after treatment.** iRBCs were treated with 2 µM UA2239 starting at 6 hpi or 36 hpi or with 1.5 µM C2 starting at 36 hpi. Parasites were processed for analysis by electron microcopy at 44-46 hpi.
